# Pan-cancer whole genome comparison of primary and metastatic solid tumors

**DOI:** 10.1101/2022.06.17.496528

**Authors:** Francisco Martínez-Jiménez, Ali Movasati, Sascha Brunner, Luan Nguyen, Peter Priestley, Edwin Cuppen, Arne Van Hoeck

## Abstract

Metastatic cancer remains almost inevitably a lethal disease. A better understanding of disease progression and response to therapies therefore remains of utmost importance. Here, we characterize the genomic differences between early-stage untreated primary tumors and late-stage treated metastatic tumors using a harmonized pan-cancer (re-)analysis of 7,152 whole-genome-sequenced tumors. In general, our analysis shows that metastatic tumors have a low intra-tumor heterogeneity, high genomic instability and increased frequency of structural variants with comparatively a modest increase in the number of small genetic variants. However, these differences are cancer type specific and are heavily impacted by the exposure to cancer therapies. Five cancer types, namely breast, prostate, thyroid, kidney clear carcinoma and pancreatic neuroendocrine, are a clear exception to the rule, displaying an extensive transformation of their genomic landscape in advanced stages. These changes were supported by increased genomic instability and involved substantial differences in tumor mutation burden, clock-based molecular signatures and the landscape of driver alterations as well as a pervasive increase in structural variant burden. The majority of cancer types had either moderate genomic differences (e.g., cervical and colorectal cancers) or highly consistent genomic portraits (e.g., ovarian cancer and skin melanoma) when comparing early- and late-stage disease. Exposure to treatment further scars the tumor genome and introduces an evolutionary bottleneck that selects for known therapy-resistant drivers in approximately half of treated patients. Our data showcases the potential of whole-genome analysis to understand tumor evolution and provides a valuable resource to further investigate the biological basis of cancer and resistance to cancer therapies.

## Introduction

Metastatic spread involves tumor cells detachment from a primary tumor, colonization of a secondary tissue, and growth in a hostile environment^1,2^. Advanced metastatic tumors are frequently able to resist aggressive treatment regimes^3^. Despite the many efforts to understand these phenomena^4–8^, we still have limited knowledge of the contribution of genomic changes that equip tumors with these extraordinary capacities. Thus, it is essential to characterize genomic differences between primary and metastatic cancers and quantify their impact on therapy resistance to be able to understand and harness therapeutic interventions that establish more effective and more personalized therapies^9^.

Multiple large-scale genome-sequencing efforts have been devoted to profiling the genomic landscape of primary tumorigenesis by relying on clinical panels^10^ or on whole-exome sequencing^11^ or whole-genome sequencing^12,13^ strategies. Similarly, recent efforts have characterized large cohorts of patients with metastatic tumors^14,15^. Several cancer type specific projects have also reported analysis of primary and metastatic matched biopsies in urothelial carcinomas^16^, breast cancer^17^, kidney clear cell carcinoma^18^ and prostate carcinoma^19,20^, among others^21^. However, the ethical and logistical challenges associated with the collection of paired biopsy data hampers the extrapolation to thousands of patients with multiple cancer types.

To circumvent this issue, most large-scale comparisons between primary and metastatic tumors have relied on unmatched whole-exome data or have adopted more targeted approaches with a specific focus on driver gene landscapes^22–24^. These efforts have frequently involved separated processing pipelines for primary and metastatic cohorts, complicating the analysis of genomic features that are highly sensitive to the selected data-processing strategy^25,26^. An impressive recent study that uniformly analyzed more than 25,000 tumors^27^ has provided a comprehensive overview of the genomic differences, driver alteration patterns and organotropism using clinical gene-panel sequencing as a base. However, this genomic analysis approach, which is limited to several hundreds of genes, prevented the exploration of the full spectrum of genomic alterations that play a role in tumorigenesis, such as structural variation and mutational scarring caused by intrinsic and extrinsic forces.

Here, we present a large-scale unified analysis of more than 7,000 whole-genome sequenced (WGS) paired tumor-normal samples (re-)analyzed by the same data-processing pipeline. This dataset enabled cancer type specific comparisons of whole-genome features in 22 cancer types with high representation in both primary and metastatic patients. We investigated differences in tumor clonality, genomic instability markers, whole-genome-duplication (WGD) rates, tumor mutation and structural variant (SV) burden and clock-based molecular signatures and assessed the contribution of cancer treatments to the observed differences. We also explored disparities in the driver gene landscape and their implications for therapeutic actionability. Finally, we identified known and new associations between certain driver alterations and exposure to various treatments. The harmonized genomic dataset used in our study constitutes the largest repository of uniformly analyzed cancer WGS data and is publicly available to the scientific community to better understand the full spectrum of DNA alterations as well as to study their role in tumor evolution and response to therapy.

## Results

### 7,152 uniformly processed whole-genome-sequenced paired tumor-normal samples from patients with primary and metastatic tumors

To characterize the genomic differences between primary and metastatic tumors, we created a uniformly analyzed dataset of matched tumor and normal genomes from patients with primary and metastatic cancer. We first collated the Hartwig Medical Foundation (Hartwig) dataset, which includes 4,784 samples from 4,375 metastatic patients (from which 2,520 samples were previously described^14^). Then, we re-processed 2,835 primary tumor samples from the Pan-Cancer Analysis of Whole Genomes (PCAWG) consortium using the open-source Hartwig analytical pipeline^14,28^ to harmonize somatic calling, standardize broad functional annotations of events and eliminate biases caused by processing pipelines and filter conditions (Supp. Fig. 1, Supp. Table 1). Reassuringly, per-sample comparison of the number of single base substitutions (SBSs), double base substitutions (DBSs), indels (IDs), and SVs revealed a strong agreement between our results and the consensus calls originally generated by the PCAWG consortium (see Supp. Note 1). Additionally, our processing pipeline strategy was minimally affected by differences in sequencing coverage, enabling a reasonable comparison of WGS samples from heterogeneous sources (decreased sensitivity at 60× and 38× coverage, typical for PCAWG samples, compared with 109× coverage, typical for Hartwig samples between 0% and 5% for simple mutations and between 8% and 14% for SVs respectively) (Supp. Note 1). A total of 7,152 tumor samples from 58 cancer types met the processing pipeline quality standards (see methods and Supp. Fig. 1a) and constitutes one of the largest publicly available datasets of WGS primary and metastatic tumors. The flexible nature of the processing pipeline, which does not require extensive parameter fine-tuning, enables its application to other whole-genome-sequencing projects, thereby providing an excellent opportunity for future integrative analyses that incorporate this dataset.

To explore genomic differences between primary and metastatic tumors, we focused on 22 cancer types from 14 tissues with sufficient sample representation (i.e., at least 15 unique patients in both the primary and metastatic cohorts), which totaled 5,751 tumor samples (1,916 primary and 3,835 metastatic) (Fig. 1a, Supp Fig 1a). Within this dataset, patients with metastatic tumors were slightly older at biopsy than patients with primary tumors (mean of 2.09 years older across all cancer types). In particular, the mean age at biopsy in primary and metastatic cohorts, respectively, was 59.9 and 67.9 years in patients with prostate carcinoma, 52.8 and 63.6 years in patients with thyroid carcinoma, and 49.6 and 70.3 years in patients with diffuse B cell lymphoma. Consistent gender proportions were observed across all cancer types except for thyroid adenocarcinomas, which had higher male representation in the metastatic cohort (metastatic: 72% male, 28% female; primary: 25% male, 75% female). Treatment information was available for 53% metastatic patients. This information is essential to gauge the contribution of this evolutionary bottleneck to the genomic differences between primary and metastatic tumors (Fig. 1a). Finally, biopsy location was explicitly annotated in 82.4% of metastatic patients, including 12.6% biopsies from metastatic lesions in the primary tissue (local), 16% in lymph nodes and 53.8% in distant locations. Biopsy locations were highly tumor type specific and likely reflected both the dissemination patterns of the tumors and the challenges associated with collection of clinical samples. In summary, we generated a harmonized dataset of 7,152 primary and metastatic WGS tumor samples from 58 cancer types including 22 cancer types with sufficient representation in both early and late clinical stages to allow for a systematic comparison (Supp. Table 1).

**Figure 1.**
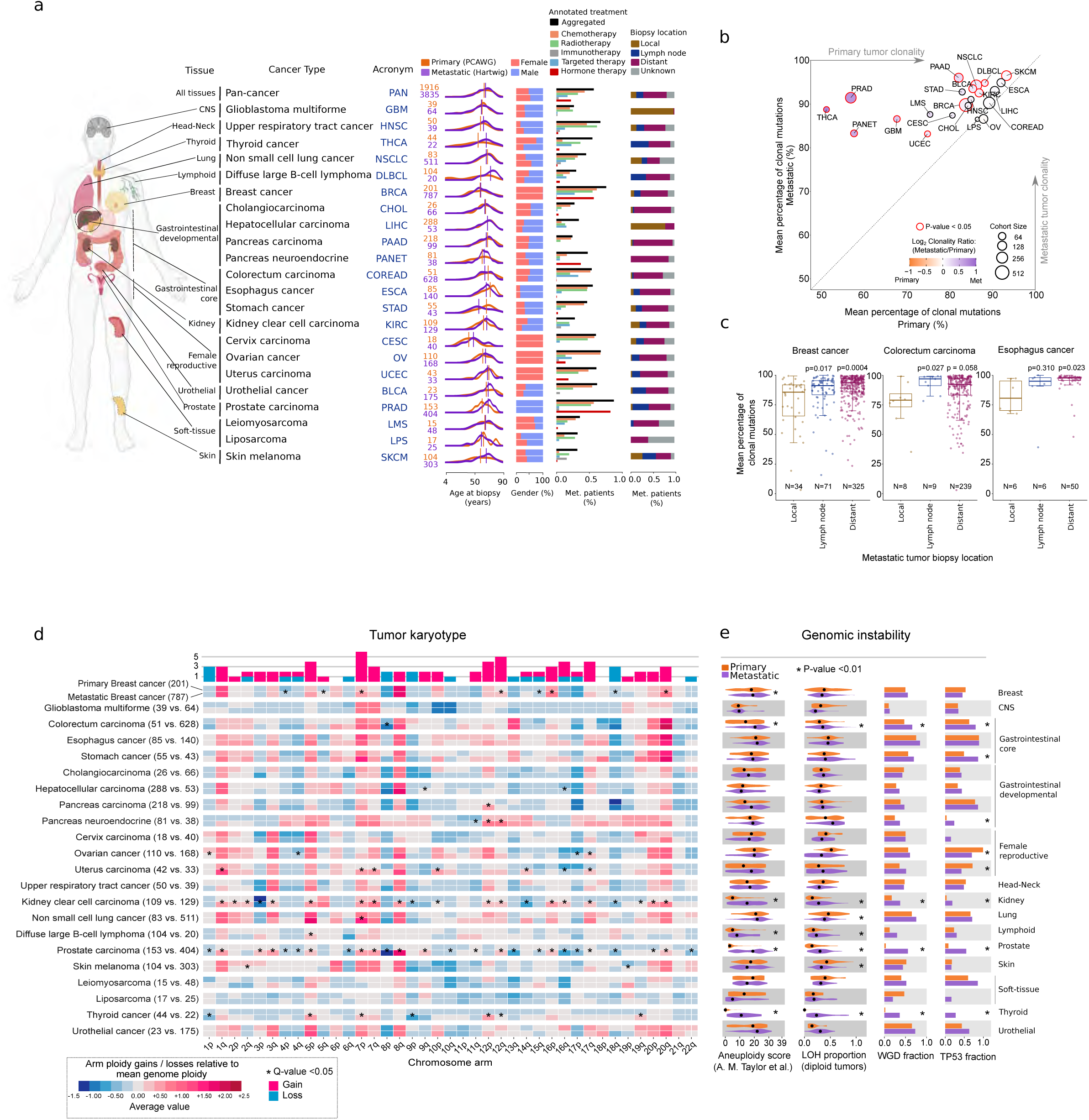
Overview and global genomic features of primary and metastatic tumors. **a)** Anatomic location of the 22 cancer types included in this study. Cancer types are ordered according to their tissue of origin. For each cancer type the following information is represented, from left to right: cancer type acronym, number of samples in primary and metastatic dataset (top and bottom, respectively), age at biopsy (in years), gender, type of treatment of metastatic patients and biopsy site from metastatic tumors. Partially created with BioRender.com. **b)** Mean percentage of clonal mutation in primary (x-axis) and metastatic (y-axis) tumors. Dots are coloured according to Log_2_ of the clonality ratio (metastatic divided by primary). Size of dots are proportional to the total number of samples (primary and metastatic). Red edge lines represent a Mann-Whitney p-value < 0.05. **c)** Tumor clonality according to the metastatic biopsy location in breast, colorectal and esophagus cancer (from left to right). N, number of samples in the group. p, Mann-Whitney p-value if p-value < 0.05. ns, not significant. Box-plots: center line, median; box limits, first and third quartiles; whiskers, lowest/highest data points at first quartile minus/plus 1.5× IQR. **d)** Tumor karyotype. Heatmap representing the mean chromosome arm ploidy gain and losses relative to the mean genome ploidy in primary (top) and metastatic (bottom) tumors. Asterisks represent significantly different mean distributions between primary and metastatic tumors (Mann-Whitney adjusted p-value <0.05). Top bars represent the cumulative number of significant gain/losses in the metastatic cohort compared to the primary. **e)** Comparison of four genomic features between primary (top) and metastatic tumors (bottom). From left to right, aneuploidy score from ref^31^, proportion of genome undergoing LOH in diploid samples, fraction of samples bearing whole genome duplication (WGD) and TP53 alterations. Black dots represent the median values. Asterisks represent Fisher’s exact test p-value < 0.01 for discrete features (WGD and TP53) and Mann-Whitney p-value < 0.01 for the continuous features.

### Global genomic characteristics of primary and metastatic tumors

We first explored global genomic differences between primary and metastatic tumors across the aforementioned 22 cancer types. Metastatic tumors showed an overall increase in clonality compared with their primary tumor counterparts (Fig. 1b). Particularly, 12 cancer types had a significantly higher metastatic average clonality ratio, ranging from 3.2% increased mean clonality in skin melanoma to 37% increased mean clonality in thyroid carcinoma. Interestingly, within the group of patients with metastatic breast cancer, distant and lymph node tumor biopsies showed significantly higher clonality ratios compared with local metastatic lesions (Fig. 1c). This increase in clonality was also observed in distant tumor biopsies of esophagus cancer and colorectal carcinomas (Fig. 1c). Nevertheless, the biopsy location did not influence tumor clonality in other cancer types such as non-small cell cancer and skin melanoma (Supp. Fig 1b), suggesting that patterns of tumor dissemination are highly tumor type specific^8^. Thus, our results thus support the notion that metastatic lesions generally have lower intra-tumor heterogeneity^27^, which may be explained by a single major subclone seeding event from the primary cancer and/or by severe evolutionary constraints imposed by anti-cancer therapies.

Karyotype comparison revealed a generally conserved portrait, which was strongly shaped by the tissue of origin^29^, and where only two cancer types showed substantial karyotypic changes (kidney clear cell and prostate carcinomas, >10 chromosome arm gains/losses) and three additional cancer types displayed moderate changes (breast, uterus, and thyroid carcinomas, >5 chromosome arm gain/losses) in the metastatic cohort compared with the primary cohort (Fig. 1d, Supp. Table 2). The majority (63 of 84) of significant discrepancies were associated with an increased frequency of chromosomal arm gains in the metastatic dataset, while only 21 discrepancies were caused by an increased frequency of chromosome arm losses. Chromosome arms 7p, 12p, 5p and 17q, which are relatively enriched in oncogenes^30^, were the most recurrent chromosome arm gains. Conversely, chromosome arms 1p and 18q, highly enriched in tumor suppressor genes^30^, were the most recurrent losses in the metastatic cohort (Fig. 1d).

We next investigated differences in four well-studied genomic instability markers: chromosomal aneuploidy score^31^, loss of heterozygosity (LOH) genome fraction in diploid tumors, WGD^32^, and *TP53* alterations^32,33^ (Fig. 1e, Supp. Table 2). Four cancer types (i.e., colorectal, kidney clear cell, prostate, and thyroid) showed persistent increases in the four genomic instability markers in the metastatic cohort, whereas four additional cancer types (i.e., breast, stomach, pancreas neuroendocrine and diffuse B cell lymphoma) had some form of increased genomic instability in the metastatic cohort. Our results thus confirmed that genomic instability is a hallmark of advanced tumors^4,9,27,29,34,35^ and revealed that the majority of cancer types have already acquired variable degrees of this genomic feature early in tumor evolution. However, certain cancer types, such as prostate, kidney, and thyroid, significantly increased the level of genomic instability in later evolutionary stages, which were, in turn, associated with substantial karyotypic changes.

### Tumor mutation burden

The unified processing of both primary and metastatic tumor samples enables quantitative and qualitative comparison of SBSs, DBSs and ID burden, which we refer to collectively as tumor mutation burden (TMB). We found that the TMB in metastatic tumors was only moderately increased compared with primary tumors across the 22 cancer types tested (fold-change increases of 1.22 ± 0.48 for SBS, 1.52 ± 0.85 for DBS and 1.43 ± 0.55 for IDs; mean ± standard deviation [SD]). In fact, more than 60% of the cancer types (13 of 22) had no significant increase in mutation burden for any mutation type. Only five cancer types (breast, cervical, thyroid, prostate carcinoma, and pancreas neuroendocrine) had a consistent increase for the three mutation types at the metastatic stage, although the mutation profiles lacked systematic differences between primary and metastatic tumors (Fig. 2b, Supp Fig. 2a). These results show that TMB is not necessarily indicative of tumor progression status and that the mutational spectra are tightly shaped by the mutational processes that were already active before and during primary tumor development.

**Figure 2.**
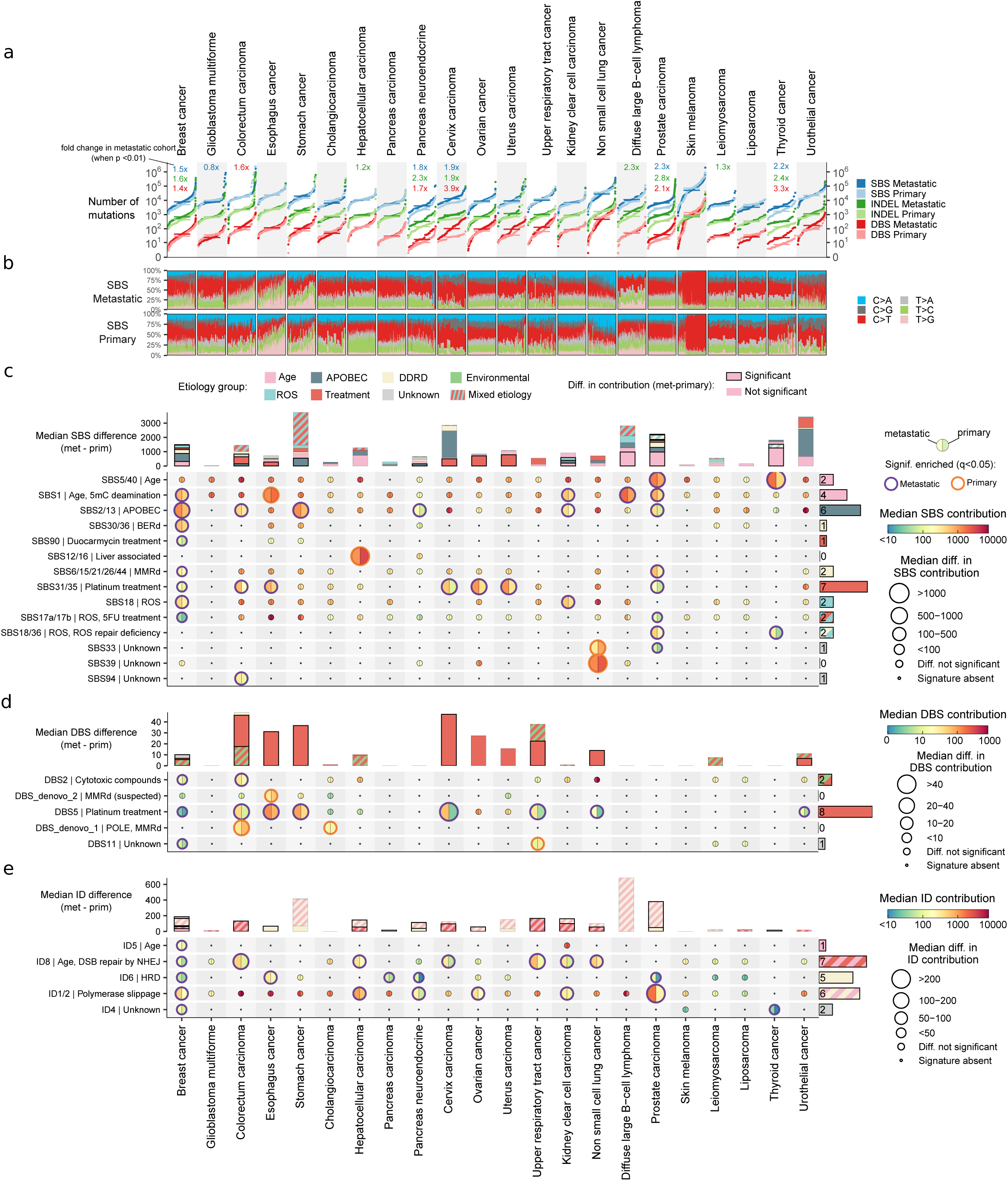
Tumor mutation burden and mutational processes. **a)** Cumulative distribution function plot (samples were ranked independently for each variant type) of tumor mutation burden for each cancer type for SBS (blue), IDs (green) and DBS (red). Horizontal lines represent median values. Fold change labels are included only when Mann-Whitney comparison rendered a significant p-value < 0.01. **b)** SBS Mutational spectra of metastatic (top) and primary (bottom) patients. Patients are ordered according to their TMB burden. **c)** Main panel, moon plot representing the mutational burden differences attributed to each mutational signature in metastatic (left) and primary (right). Edge thickness and colors represent significant differences (q<0.05, ±1.4x fold change) and the direction of the enrichment, respectively. The size of circles are proportionate to the mutation burden difference. Right bars, number of metastatic cancer types with a mutational signature significant enrichment. Top stacked bars represent the cumulative signature exposure difference. Thicker bar edge lines represent significance. Bars are coloured according to the annotated etiology. Only mutational signatures with known etiology or with at least one cancer type with significant metastatic enrichment are included. **d)** and **e)** equivalent for DBS and IDs, respectively. Diff., difference. Muts. mutations. Sig., mutational signature. Mut. mutational. Susp., suspected.

### Mutational processes in primary and metastatic tumors

To assess whether the TMB differences may be attributed to differential activity of environmental or endogenous mutational processes, we conducted a tissue type specific mutational signature *de novo* extraction that resulted in representative mutational signatures of 69 SBSs, 12 DBSs, and 19 IDs (Supp. Table 3). Most of these (50 of 69 SBSs, 8 of 12 DBSs, and 12 of 19 IDs) mapped onto the well-described mutational signatures in human cancer^36^. Moreover, we inferred the suspected etiology for two novel DBS mutational signatures associated with mismatch repair deficiency (MMRd) (DBS_denovo_2) and MMRd/POLE hypermutation (DBS_denovo_1) (Supp. Table 3).

By comparing the activities of all mutational processes, we found that mutations caused by cytotoxic treatments were enriched in 11 cancer types (Fig. 2c-e). Platinum-based chemotherapies (SBS31/SBS35 and DBS5) showed the strongest mutagenic effect with 429 ± 266 (mean ± SD) SBS mutations and 23 ± 15 (mean ±SD) DBS mutations on average per sample. In fact, the excess in DBS mutation burden observed in six cancer types (stomach, esophagus, cervix, upper respiratory tract, non-small cell lung, and urothelial cancer) could be fully attributed to platinum treatment (Fig. 2d top bars). Likewise, the radiotherapy ID signature^37^ (ID8) was systematically enriched in multiple cancer types as a response to widespread exposure to radiation-based treatment, whereas the 5-fluorouracil^38,39^ (SBS17a/b), duocarmycin^40^ (SBS90) and polycyclic aromatic hydrocarbon (PAH) metabolites from chemotreatments^41^ (DBS2) occasionally led to a greater mutation burden in metastatic tumors in a tumor type specific manner (Fig. 2c-e).

The systematic enrichment of SBS2/13 mutations in metastatic cancers suggests enhanced activity of APOBEC mutagenesis during the progression of advanced tumors. Specifically, our results revealed an increase in APOBEC mutation burden of 316 ± 173 (mean ± SD) mutations per sample in six metastatic tumors (breast, colorectal, stomach, kidney, prostate, and pancreas neuroendocrine) that reached statistical significance, being breast and stomach cancers the ones with the strongest increase (>500 APOBEC mutations per sample in both cancer types). Other cancer types, such as cervical and urothelial cancers, also showed enhanced APOBEC activity (>1800 mutations per sample), but they did not reach significance owing to high intrinsic APOBEC activity in the primary tumors. The metastatic breast cancer samples also had a higher percentage of APOBEC hypermutation than primary tumors (14% vs 5%, Supp. Fig. 2c, Supp. Table 3).

Five metastatic cancer types also displayed more mutations from the clock-like mutational processes, including four cancer types (breast, prostate, diffuse B-cell lymphoma and kidney clear cell carcinoma) that exhibited an increased SBS1 contribution and two cancer types (prostate and thyroid) that had an increased SBS5/SBS40 mutation burden. The increase in clock-like mutations in thyroid and prostate cancers, as well as diffuse B-cell lymphomas to a lesser extent, may be explained by the greater proportion of older patients with metastatic disease compared with primary disease. However, the SBS1 enrichment was also present in cancer types in the primary and metastatic cohorts with highly similar age population distributions (see Fig. 1a). Thus, our results revealed an increase in SBS1 mutations in advanced tumor stages that cannot be explained by older patients’ ages at the time of biopsy.

Additional more-focused analyses may achieve a better understanding of the observed mutational signatures with differing activity in a small subset of cancer types (such as oxidative stress-induced and DNA repair related mutations) and those lacking mechanistic etiology warrants additional more focused analyses. All contributions can be found in Supp. Table 3.

### Differential SBS1 mutation rates in primary and metastatic cancers

To investigate the apparent contradiction of increased SBS1 mutation burden in the metastatic cohort, despite primary and metastatic cohorts having a similar age at biopsy, we assessed their SBS1 mutation burden by the age of biopsy separately for both cohorts (see methods). As expected, rates of SBS1 mutation acquisition were highly tissue specific^42,43^, and SBS1 mutation burden increased linearly with age in the majority of cancer types in both primary and metastatic cohorts (Pearson’s R > 0.1, 17 of 22 tumor types, Supp. Fig. 3a, Supp. Table 4). However, four cancer types (i.e., breast, prostate, kidney clear cell, and thyroid) showed an age-independent and significant enrichment of SBS1 mutations at the metastatic stage (Fig. 3a, Supp. Fig. 3a). For instance, metastatic breast cancer had a nearly uniform fold increase of 1.46 (189 ± 17 SBS1 mutations, mean ± SD) across the ages of biopsies that was independent of tumor genome ploidy and *HER2* status (Supp. Fig. 3b-c). Importantly, this pattern was highly cancer type specific and was not observed for most cancer types, including those with similar intra-tumor heterogeneity in the primary cohort (e.g., colorectal, ovarian cancer, pancreas, and stomach cancers) (Fig. 3b, Supp. Fig. 3a). Moreover, other mutational processes that operate over the evolution of the somatic tissues (e.g., clock-like mutations attributed to SBS5/SBS40 that accumulate with age in a cell-cycle independent manner^43,44^) were not enriched (Supp. Fig. 3d).

**Figure 3.**
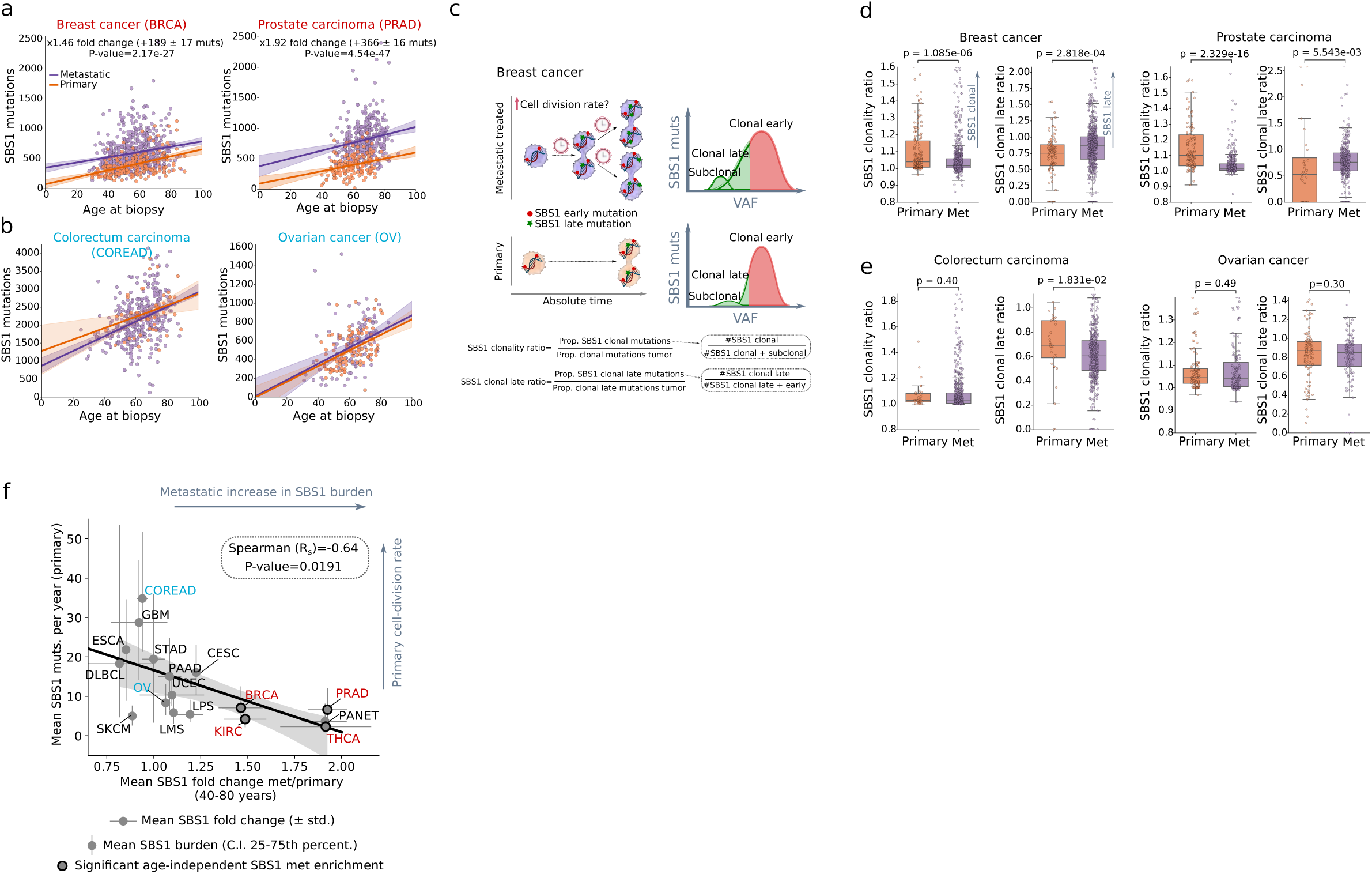
Cell cycle division rates in primary and metastatic tumors. **a)** Linear regression of the SBS1 mutation burden (y-axis) and patient’s age at biopsy (x-axis) in primary and metastatic breast (left) and prostate cancers (right) and **b)** colorectal and ovarian cancer. The mean fold change, mean SBS1 increase per year and p-value are included in cancer types with an age-independent significantly different primary and metastatic distribution. The median trendline and 99% confidence intervals of the linear regression are represented as a solid line and the adjacent shaded area, respectively. **c)** Depiction illustrating increased cell division rate in metastatic breast cancer compared to primary and its expected impact on the SBS1 variant allele frequency (VAF) distribution. Partially created BioRender.com. **d)** Comparison of global SBS1 clonality ratio and clonal late ratio between primary and metastatic in breast (left) and prostate cancers (right)**. e)** Similar for colorectal and ovarian cancers. Boxplots are defined as in Fig. 1. **f)** Spearman correlation analysis of the mean SBS1 year burden of primary tumors (y-axis) and the mean metastatic SBS1 fold change (x-axis) across the 17 cancer types with linear association between age and SBS1 accumulation. Vertical error bars represent the 25th and 75th percentile, respectively. Horizontal error bars the fold change standard deviation. Cancer types with a significantly different SBS1 mutation rate are marked by thicker marker borders. Muts, mutations.

SBS1 mutation burden has been extensively correlated with estimated stem cell division rates^45^. Therefore, an increase in age and tumor type specific SBS1 mutation burden in treated metastatic tumors may indicate that these tumors have undergone a higher number of cell divisions. However, the estimated number of years to explain the SBS1 mutation burden shift (23 and 67 years for breast and prostate cancer respectively, see Supp. Table 4) shows that this cannot be the main cause. Hence, a more plausible explanation, which also supports previous observations^46–48^, is that these metastatic tumors display accelerated cell division rates compared with their primary tumor counterparts (Fig. 3c). Supporting this hypothesis, metastatic tumors also had a lower normalized fraction of clonal SBS1 mutations as well as a greater fraction of SBS1 clonal late mutations (Fig. 3d, Supp. Fig. 3e). Of note, this pattern was not observed in cancer types with consistently high SBS1 mutagenic dynamics (Fig. 3e) and was indistinguishable for SBS5/SBS40 mutations (Supp. Fig. 3f).

To understand the underlying factors that lead to the increased proliferation rates in some cancer types and not in others, we explored the relationship between the yearly rate of SBS1 mutation accumulation in primary tumors (a proxy of stem cell division rates^45^) and the estimated fold change of the SBS1 mutation rate in the metastatic cohort. Remarkably, we observed a strong negative association between these two factors (Spearman R_s_= -0.64, p-value = 0.01, Fig. 3f), which was consistent when relying on independent measurements of primary tumor turnover rates (Supp. Fig. 3h, Supp. Table 4). This indicates that tumors with an intrinsically active turnover rate (e.g., colorectal and ovarian cancers) preserve their high proliferation rates, while others with low cell division rates (e.g., breast, prostate, kidney, and thyroid cancers), may acquire higher proliferation rates during the course of cancer progression and treatment exposure.

### Structural variant burden

The total number of SVs per tumor revealed an extensive increase in SV burden in the metastatic tumors (fold change 1.9 ± 1.2, mean ± SD). In fact, 14 of 22 (64%) cancer types showed a significant median increase in total SV burden in the metastatic tumors (Fig. 4a, Supp. Table 5), which cannot be explained by differences in sequencing coverage or tumor clonality (Supp. Note 1). Importantly, these cancer types not only included the eight cancer types with increased genomic instability markers, but also six additional cancer types (i.e., esophagus, hepatocellular, pancreas carcinoma, cervix, non-small cell lung cancer, and urothelial cancer) (Fig. 4a).

**Figure 4.**
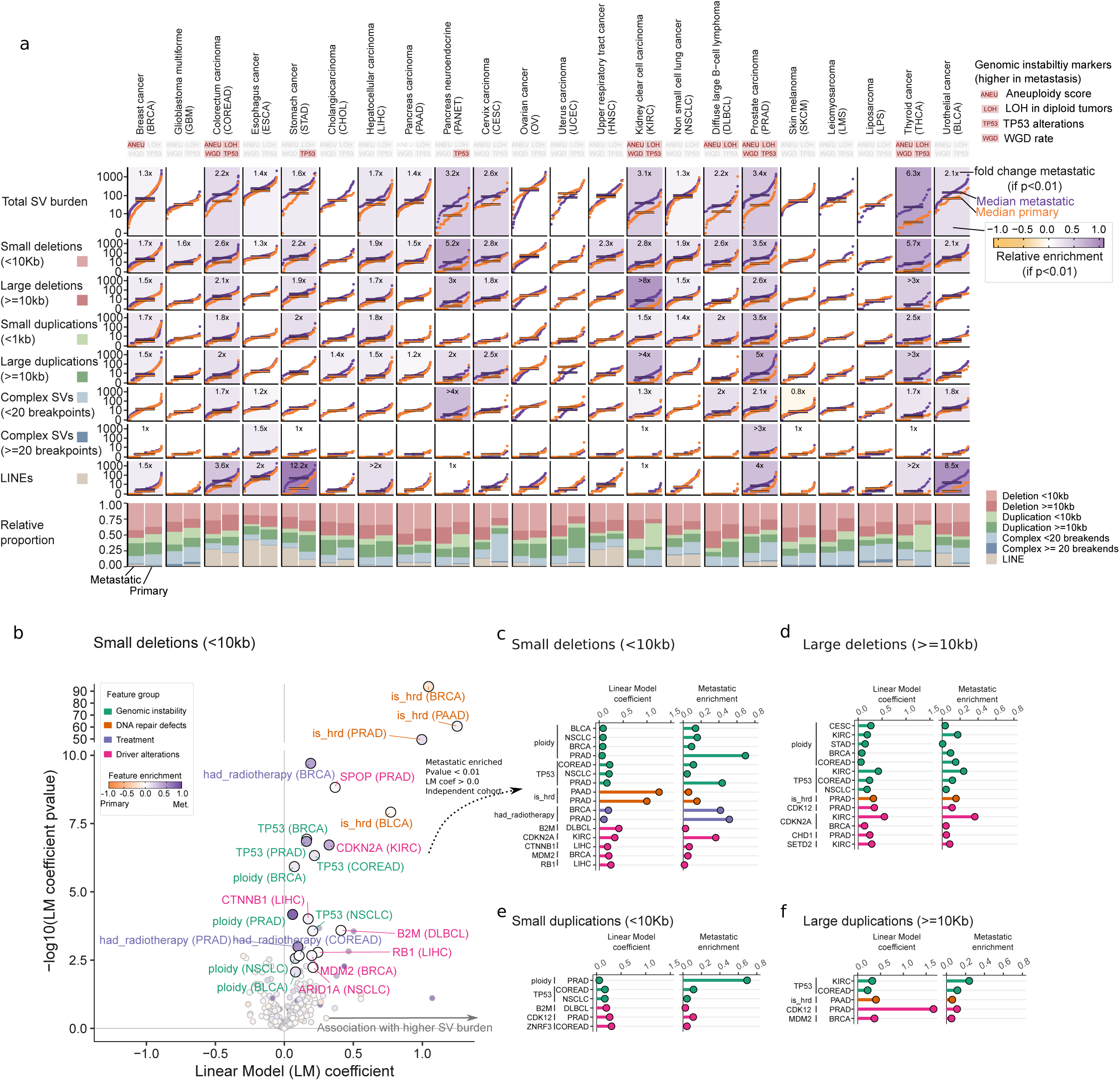
Structural variant burden and associated genomic features. **a)** Top rectangles represent the four genomic instability features defined in Fig. 1e. A red background represents significant enrichment in the metastatic cohort. S-plots, cumulative distribution function plot (samples ranked independently for each SV type) of tumor mutation burden for each cancer type for (from top to the bottom) the aggregated structural variant (SV) burden, small deletions (<10kb), large deletions (>=10kb), small duplications (<10kb), large duplications (>=10kb), complex events (<20 breakpoints), complex events (>=20 breakpoints) and LINEs insertions. Horizontal lines represent median values. Backgrounds are coloured according to the relative enrichment, defined as: log_10_(median SV type burden in metastatic tumors + 1) − log_10_(median SV type burden in primary tumors + 1). Fold change labels and coloured backgrounds are displayed when Mann-Whitney comparison renders a significant p-value < 0.01. Fold change labels are displayed with ‘>’ when the SV burden for primary tumors is 0 (see methods for more details). For each cancer type, bottom bar plots represent the relative fraction of each SV type in the metastatic (left) and primary (right) datasets. **b)** Volcano plot representing the cancer type specific regression coefficients (x-axis) and significance (y-axis) of clinical and genomic features against the number of small deletions. Each dot represents one feature in one cancer type. Labels are coloured according to the feature category. Dots are coloured by the frequency enrichment in metastatic (purple) or primary (orange) patients. LM, linear model. Coef, coefficient. **c)** Lollipop plots representing the regression coefficients (left, relative to panel b. x-axis) and metastatic enrichment (right, relative to dots color from panel b.) of features associated with small deletions. Only significant features (LM>0.0, p-value < 0.01 and with independent significance in primary or metastatic tumors) enriched in metastatic patients (enrichment > 0.0) are displayed. **d), e)** and **f)** are identical but referring to large deletions, small duplications and large duplications, respectively.

Breaking down the comparison into specific SV types revealed that small (<10kb) deletions, which also contributed most to the global SV burden, showed a strong metastatic enrichment (2.15 ± 1.3 fold change in 16 of 22 cancer types with significant enrichment, mean ± SD, Fig. 4a, Supp. Fig. 4a). Moreover, larger (≥10kb) deletions and duplications had a similar pan-cancer enrichment, although with slightly lower fold changes that varied from 1.5 up to 1.9. Moreover, complex SVs with ≥20 breakpoints, encompassing events such as chromothripsis and chromoplexy, were particularly enriched in esophagus cancer (1.5-fold) and prostate cancer (3-fold). Finally, a strong cancer type specific metastatic enrichment was also noted for the long interspersed nuclear element (LINE) insertions, with an increased fold change of 12.2 and 8.5 in stomach and urothelial cancer, respectively.

Compared with TMB, the SV analyses revealed a much more widespread effect, with larger increases per metastatic cancer type that affected almost every cancer type studied, indicating that metastatic tumors appear to evolve primarily by genomic changes at the structural level.

### Genomic and clinical features associated with structural variant burden

We next sought to unravel the underlying features associated with the observed increase in SV burden in metastatic tumors using linear regression models (Fig. 4b-f, Supp. Fig. 4b-f, Supp. Table 5). Our approach confirmed previously described cancer type specific driver-induced SV phenotypes, including HRd (*BRCA1/2*)^49,50^, *CDK12*^51^, and *MDM2*^52^, which are more frequently mutated in metastatic pancreatic, prostate, and breast carcinomas, respectively. Reciprocal duplications induced by *CDK12* and *CCNE1* alterations^53^ likely explain the enrichment of complex rearrangements in prostate and esophagus cancer, respectively. We also found potential novel associations, such as large deletions linked to the chromatin regulators *SETD2* and *CHD1*. Finally, the cell cycle checkpoint *CDKN2A* showed a pervasive association with deletions in breast and kidney clear cell carcinomas (Fig. 4b-d, Supp. Fig. 4b). However, this may not necessarily imply causation, because the tumor suppressor *CDKN2A* is frequently inactivated in tumors in which deletion signatures are common^54^.

We also found that genomic instability features (i.e., genome ploidy and *TP53* alterations) showed a strong pan-cancer association with deletions and, to a lesser extent, with duplications (Fig. 4b-f, Supp. Fig. 4b-f), supporting the established role of WGD^32^ and *TP53*^31^ loss in the ubiquitous generation of large genomic alterations. As mentioned earlier (see Fig. 1e), these features were generally more prevalent in patients with metastatic tumors and thus very likely contributed to the observed SV increase in metastatic tumors.

Finally, prior exposure to radiotherapy treatment was strongly associated with small deletions in breast and prostate cancers (in agreement with radiotherapy-treated secondary malignancies^55^, gliomas^37^, and healthy tissue^56^). This suggests that, among common anti-cancer therapies, radiotherapy in particular significantly contributes to the SV landscape in treatment-surviving cancer cells.

### Cancer driver gene alterations in primary and metastatic tumors

Comparison of the total number of driver gene alterations per tumor sample revealed a moderate increase in the metastatic cohort (a mean of 4.5 and 5.2 driver alterations per sample in primary and metastatic tumors, respectively), including 11 (50%) tumor types with a significant increase (Fig. 5a). Prostate adenocarcinoma (3.02 driver alterations per sample), pancreas neuroendocrine (2.16), thyroid carcinoma (1.7), and kidney clear cell carcinoma (1.63) showed the strongest increases (>1.5 driver alterations per patient), whereas the majority of cancer types showed a mean increase below 1.5 driver alterations per sample. All mutation types (amplifications, deletions, and mutations) tended to show consistent biases toward increases in metastatic tumors (Fig. 5a), with small variant mutations contributing the most (2.3 and 2.6 driver mutations per sample in primary and metastatic tumors, respectively).

**Figure 5.**
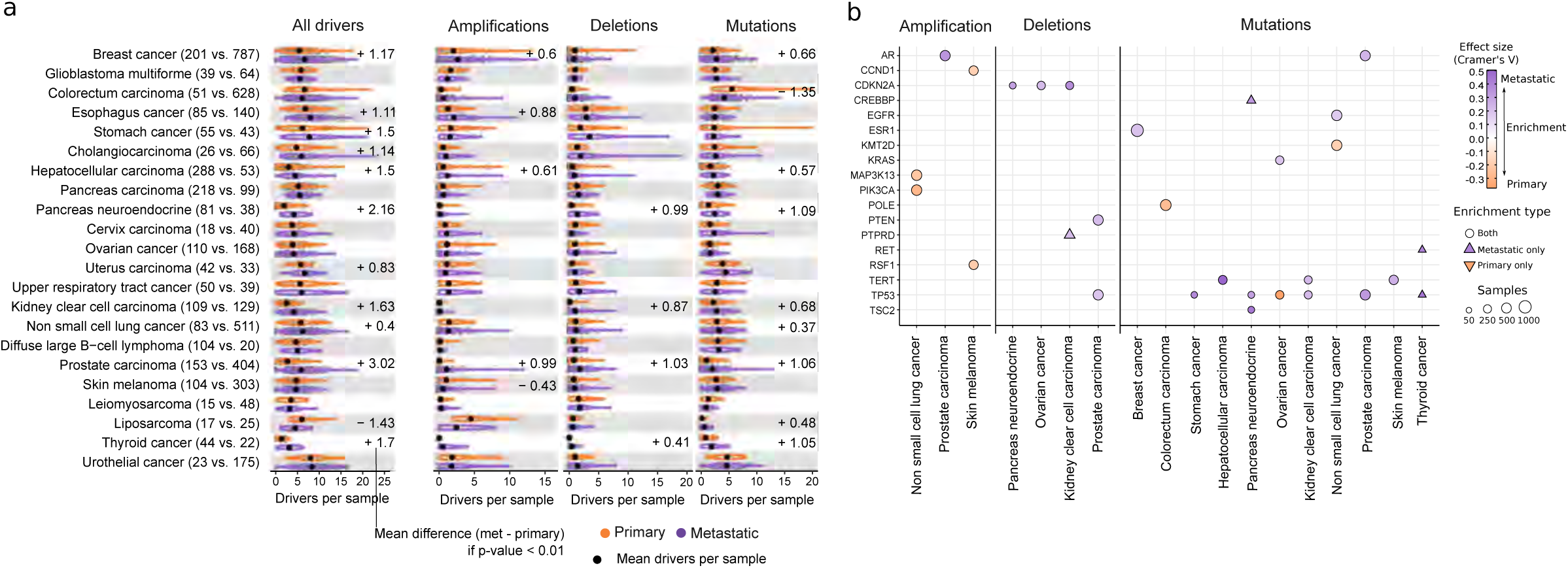
Driver alterations in primary and metastatic tumors. **a)** Cancer type specific distribution of number of driver alterations, amplifications, deletions and mutations per patient in primary (top) and metastatic (bottom). Black dots represent the mean values. Labels display mean differences (metastatic - primary) in cancer types with a significant difference. **b)** Heatmap representing the cancer genes displaying significant mutation frequency differences between primary and metastatic tumors. Circles denote mutation frequency enrichment in both cohorts while triangles facing upwards and downwards represent drivers that are exclusively enriched in metastatic and primary cohorts, respectively. Marker size is relative to the total number of mutated samples. Colors represent the direction of the enrichment.

Comparison of gene- and cancer-type frequencies revealed that only 18 genes had a significant frequency bias in at least one cancer type (29 gene and cancer-type pairs in total, Fig. 5b, Supp. Fig. 5a-b, Supp. Table 6). The majority (21 of 29, 72%) of the significant pairs had enrichment toward higher metastatic frequency, including four driver genes that were exclusively mutated in metastatic tumors (*PTPRD* in kidney clear cell carcinoma, *CREBBP* in pancreas neuroendocrine, and *RET* and *TP53* alterations in thyroid carcinoma). Moreover, most metastatic-enriched cancer drivers had a cancer type specific enrichment that included well-established resistance gene drivers associated with anti-cancer therapies, such as *AR* and *ESR1* alterations in patients with prostate and breast cancer treated with hormone deprivation therapies^57,58^, and *EGFR* mutations in patients with metastatic non-small cell lung cancer often treated with anti-EGFR inhibitors. Nevertheless, three driver genes (i.e., *TP53*, *CDKN2A,* and *TERT*) showed a metastatic enrichment across multiple cancer types (Fig. 5b), indicating that alterations of these genes may enhance aggressiveness by disturbing pan-cancer hallmarks of tumorigenesis. In fact, TP53 alterations have been extensively linked to genomic instability^32,33^, while *CDK2NA* and *TERT* are key regulators of cell proliferation, two pathways that are often perturbed in metastatic tumors^59^ (see earlier). Finally, some driver genes were strongly enriched in primary tumors, such as *KMT2D* mutations in non-small cell lung cancer (primary 13.7%, and metastatic 2.1%) and *POLE* mutations in primary colorectal carcinomas (primary 13%, metastatic 0.5%). Of note, the higher prevalence of *POLE* mutations in primary tumors supports the increased SBS10a/b exposure (i.e., *POLE* hypermutation) in primary colorectal cancer (see Fig. 2d). Whether these alterations are indicators of better prognosis or drivers of subclonal expansion warrants further investigation.

We next investigated whether the reported driver differences may impact on potential clinical actionability. Cancer type specific comparison of therapeutically actionable variants revealed an overall greater fraction of patients with therapeutically actionable variants in the metastatic cohort, with high variability across cancer types (Supp. Fig. 6a, Supp. Table 7). Subsetting by A-on label variants (i.e., approved biomarkers in the specific cancer type) revealed a consistent pattern in which only breast cancer (driven by higher *PIK3CA* mutation frequency) and non-small cell lung cancer (*EGFR* and *KRAS*^G12C^ alterations) showed a significant proportional increase in the metastatic cohort (Supp. Fig. 6a-b). Evidence levels representing biomarkers in experimental clinical stages (A-off label, B-on label, and B-off label) showed a modest and tumor type dependent metastatic increase, which was mainly linked to the increased alteration frequency of *KRAS*^G12X^, *EGFR* secondary mutations, and *CDKN2A* loss in advanced tumor stages (Supp. Fig. 6b).

In conclusion, the cancer driver gene landscape is generally conserved and the observed differences are associated with hallmarks of tumor aggressiveness and resistance to anti-cancer therapies. Consequently, therapeutic options are mainly dictated by the primary tumor^60^, although advanced experimental drugs may provide relevant therapeutic opportunities for metastatic tumors in the near future.

### Treatment associated driver gene alterations

The high prevalence of resistance driver genes in late-stage tumors prompted us to devise a test that aimed to identify treatment enriched drivers (TEDs) that were either significantly enriched (i.e., treatment enriched) or exclusively found (i.e., treatment exclusive) in a cancer type and treatment specific manner (Fig. 6a and methods). Our analytical framework provided 56 TEDs associated with 24 treatment groups from eight cancer types (Fig. 6b bottom pie chart, Fig. 6c, Supp. Table 8). Of the identified TEDs, 28 of 56 (50%) were coding mutation drivers, 18 (30%) copy number amplifications, 8 (15%) non-coding drivers and 3 (5%) recurrent homozygous deletions (Fig. 6c, Supp. Fig. 7a-b). Reassuringly, the majority of the top hits were widely known treatment-resistance drivers, including *EGFR*^T790M^ mutations (Fig. 6d) and *EGFR* copy number gains in patients with non-small cell lung cancer treated with *EGFR* inhibitors^61,62^ (Supp. Fig. 7c), *AR*-activating mutations and gene amplifications in prostate cancer patients treated with androgen-deprivation therapy^58^ (Supp. Fig. 7d-e), and *ESR1*^536–538^ mutations in breast cancer patients treated with aromatase inhibitors^57^ (Supp. Fig. 7f), among others. Moreover, we also found that *TP53*, *KRAS* and *PIK3CA* alterations were recurrently associated with resistance to multiple treatments, which may indicate that these alterations are prognostic markers for enhanced tumor aggressiveness and plasticity rather than being a cancer type specific mechanism of drug resistance.

**Figure 6.**
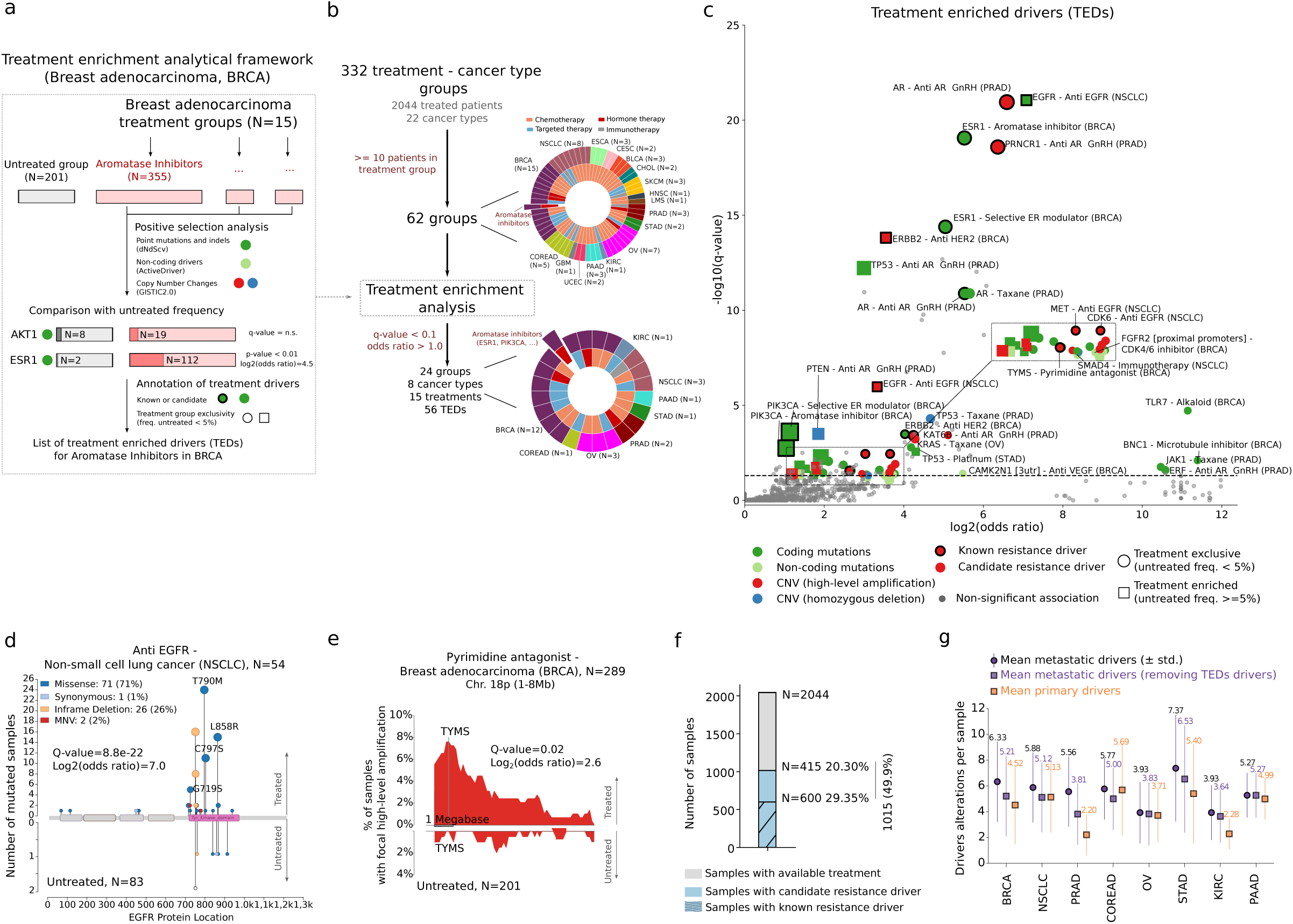
Treatment enriched drivers. **a)** Visual depiction of the analytical framework to identify treatment enriched drivers (TEDs). The example illustrates the identification of TEDs in the 355 breast cancer patients treated with aromatase inhibitors. First step, identification of cancer driver genes from coding mutations (green), non-coding mutations (soft green), copy number amplifications (red) and biallelic deletions (blue). Second, for each identified cancer driver, comparison of the mutation/copy number alteration frequency in treated and untreated patients. Third, annotation of TEDs with orthogonal evidence and type of enrichment. **b)** Left, workflow representing the number of treatment groups in each step of the analysis. Pie charts, the external layers represent the number of treatment groups analyzed (top) or with identified TEDs (bottom) coloured by cancer type. The internal layers represent the category of the corresponding treatment. The group of breast cancer patients treated with aromatase inhibitors highlighted in both charts. N, number of patients in each treatment group. **c)** Volcano plots displaying the identified TEDs. Each dot represents one cancer gene alteration type in one treatment group. X-axis displays the effect size (as log_2_[odds ratio]) and the y-axis the significance (-log_10_[q-value])). Circle markers represent TEDs exclusively mutated in the treatment group (squared makers otherwise). Markers are coloured according to the type of alteration. Known resistance drivers are denoted by thicker edgelines. **d)** Distribution of mutations along the *EGFR* protein sequence in non-small cell lung cancer patients treated with EGFR inhibitors (top) and untreated (bottom). Pfam domains are represented as rectangles. Mutations are coloured according to the consequence type. **e)** Distribution of highly-focal copy number gains in chromosome chr18p:1Mb-8Mb in breast cancer untreated patients (bottom) and treated with pyrimidine antagonists (top). TYMS genomic location is highlighted. **f)** Global proportion of metastatic treated patients with known and candidate TEDs. **g)** Mean number of driver alterations per metastatic patient before (purple circle) and after excluding TEDs (purple square) compared to primary patients (orange square). Vertical lines, standard deviation range and labels the mean number of driver alterations. BRCA, Breast cancer. CESC, Cervix carcinoma. CHOL, Cholangiocarcinoma. COREAD, Colorectal carcinoma. ESCA, Esophagus cancer. GBM, Glioblastoma multiforme. KIRC, Kidney clear cell carcinoma. LMS, Leiomyosarcoma. NSCLC, Non small cell lung cancer. OV, Ovarian cancer. PAAD, Pancreas carcinoma. PRAD, Prostate carcinoma. SKCM, Skin melanoma. STAD, Stomach cancer. THCA, Thyroid cancer. HNSC, Upper respiratory tract cancer. BLCA, Urothelial cancer. UCEC, Uterus carcinoma. MB, Megabase.

Our results also provided a long tail of candidate drivers of resistance, some of them with orthogonal evidence by independent reports (Fig. 6c, Supp. Fig. 7a-b). Examples of the latter group include *TYMS* amplification in breast cancer patients treated with pyrimidine antagonists^63^ (Fig. 6e), *PRNC1* and *MYC* co-amplifications in prostate cancer patients treated with androgen-deprivation^64^ (Supp. Fig. 7g), *SMAD4* mutations in non-small cell lung cancer treated with immunotherapy^65^, and *FGFR2* promoter mutations in breast cancer patients treated with CDK4/6 inhibitors^66^. The full TEDs catalog is provided in Supp. Table 8 and constitutes a valuable resource for investigating resistance mechanisms to common cancer therapies.

Overall, almost 50% of patients with metastatic disease with annotated treatment information harbored TEDs, including 30% with annotations of known resistance drivers and an additional 20% of patients with candidate resistance drivers derived from our analysis (Fig. 6f). We identified 0.70 ± 0.53 (mean ± SD) TEDs per metastatic sample across the eight cancer types that had reported TEDs (Fig. 6g), with prostate and breast cancers displaying the greatest prevalence of TEDs (i.e., 1.74 and 1.12 drivers per patient with prostate and breast cancer, respectively). Therefore, after excluding TEDs, primary and metastatic tumors had an approximately 40% reduction of their original differences in number of drivers per sample (from 5.2 to 4.9 mean drivers per sample in the metastatic cohort after excluding TEDs, compared with 4.5 mean drivers per sample in the primary cohort) (Fig. 6g, Supp. Table 8), indicating that an important proportion of the metastatic-enriched drivers are likely associated with resistance to anti-cancer therapies.

## Discussion

In this study we describe a cohort of >7,000 uniformly (re-)processed WGS samples from patients with primary untreated and metastatic treated tumors. Our robust analytical pipeline enabled the processing of large sets of paired tumor-normal WGS samples from diverse sequencing platforms with high efficiency and minimal human intervention. We leveraged this dataset to perform an in-depth comparison of genomic features across 22 cancer types with high representation from patients with primary and metastatic tumors.

Our analyses revealed that metastatic tumors share common genomic traits, such as high genomic instability, low intra-tumor heterogeneity, and stronger enrichment of SVs, but fewer short mutations than primary tumors. However, the magnitude of genomic differences between primary and metastatic tumors is highly cancer type specific and is strongly influenced by exposure to cancer treatments. Overall, five cancer types (prostate, thyroid, kidney clear cell, breast, and pancreas neuroendocrine cancers) showed an intense transformation of the genomic landscape in advanced tumorigenic stages. Fueled by increased genomic instability, these cancer types displayed substantial differences in TMB, clock-based molecular signatures and driver gene landscape as well as the pervasive increase in SV burden (Fig. 7a, labeled as Strong). Importantly, metastatic prostate and breast cancers regularly harbored metastatic-exclusive therapy-resistant driver gene mutations, suggesting that an important proportion of the genomic differences in metastatic tumors compared with primary tumors may be associated with clonal (re-)expansion following therapy exposure (see ref^21^ supporting this notion). However, the genomic differences in kidney, thyroid, and pancreas neuroendocrine carcinomas could not be linked to genomic markers of therapy resistance, which may indicate alternative evolutionary dynamics independent of cancer treatment. Another nine cancer types (e.g., cervix and colorectal, among others) displayed moderate genomic differences, although the global genomic portrait was conserved. Finally, the genomic landscape of the eight remaining cancer types (e.g., glioblastoma, sarcoma, and ovarian cancer, among others) was highly consistent between primary and metastatic stages, and the minimal differences were mainly attributed to exposure to cancer therapies.

**Figure 7.**
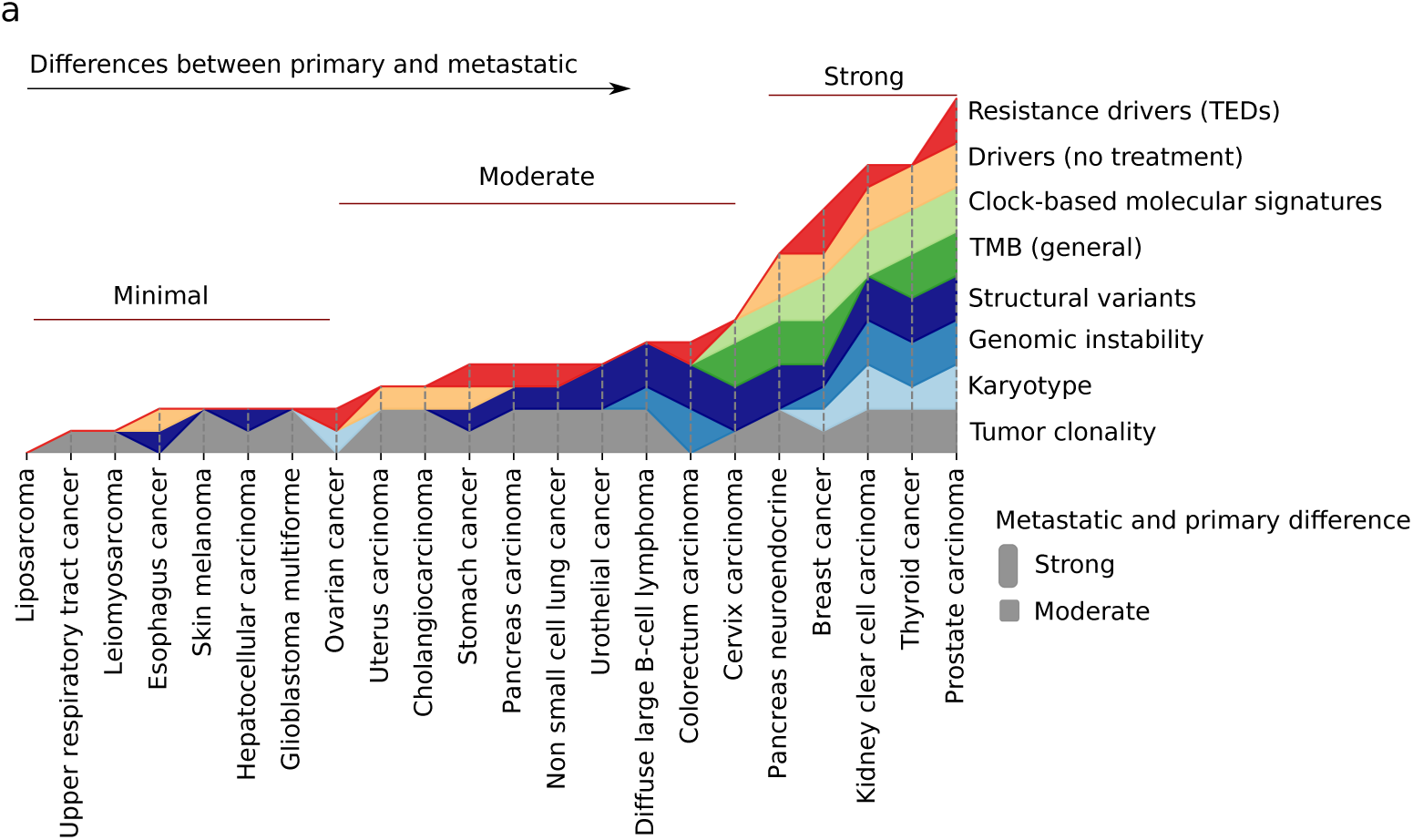
Pan-cancer differences between primary and metastatic tumors. **a)** Stacked plot representing the qualitative differences of the eight studied genomic features across the 22 cancer types included in this study. Cancer types are sorted in ascending order according to the cumulative number of diverging genomic features between primary and metastatic tumors. Each horizontal track represents a genomic feature. The presence (and height) of each feature for a specific cancer type correlates with the magnitude of the observed differences.

This study has several limitations. First, it was performed using unpaired primary and metastatic tumor biopsies. Ideally, matched biopsies from the same patient, as already implemented in cancer type specific studies^18,67,68^, would be needed to more specifically address the evolutionary dynamics of treated metastatic tumors. Moreover, the sequencing depths and tumor purity ranges used in this study are suboptimal to comprehensively profile the heterogeneous landscape of subclonal alterations, which may lead to underestimation of the extent of late-active mutational processes (e.g., treatment-induced processes). Single cell-based sequencing approaches will be instrumental to further dissect mutational landscapes independent of their clonality. In addition, the sequencing depth of the primary tumor cohort was lower and more variable than that of the metastatic tumor cohort. Although we demonstrated that this does not severely impact on the overall detectability of clonal somatic variants, individual drivers may have been occasionally missed, which may negatively impact statistical accuracy. Finally, genomic changes cannot entirely explain how tumor cells are able to colonize other organs while avoiding the strong bottlenecks imposed by the immune system^69^ or by aggressive treatment regimes. Therefore, additional information from complementary tumor *omics* studies^70,71^ and from the tumor microenvironment^72,73^ will be needed to further dissect and better understand tumor evolution and resistance to cancer therapy and eventually contribute to improved management of this deadly disease.

To conclude, our dataset constitutes a valuable resource that can be leveraged to further study other aspects of tumor evolution (as illustrated by the accompanying publication^69^ focusing on genetic immune escape alterations) as well for the development of machine learning tools to foster cancer diagnostics^74^.

## Supporting information

SuppData1_denovo_signature_contribs

SuppTable1_sample_metadata

SuppTable2_karyotype

SuppTable3_mutational_signatures

SuppTable4_ageMutationalSignatureCounts

SuppTable5_sv_load

SuppTable6_driver

SuppTable7_actionability

SuppTable8_TEDs

## Supplementary Information

### Supplementary Figures

**Supplementary Figure 1.**
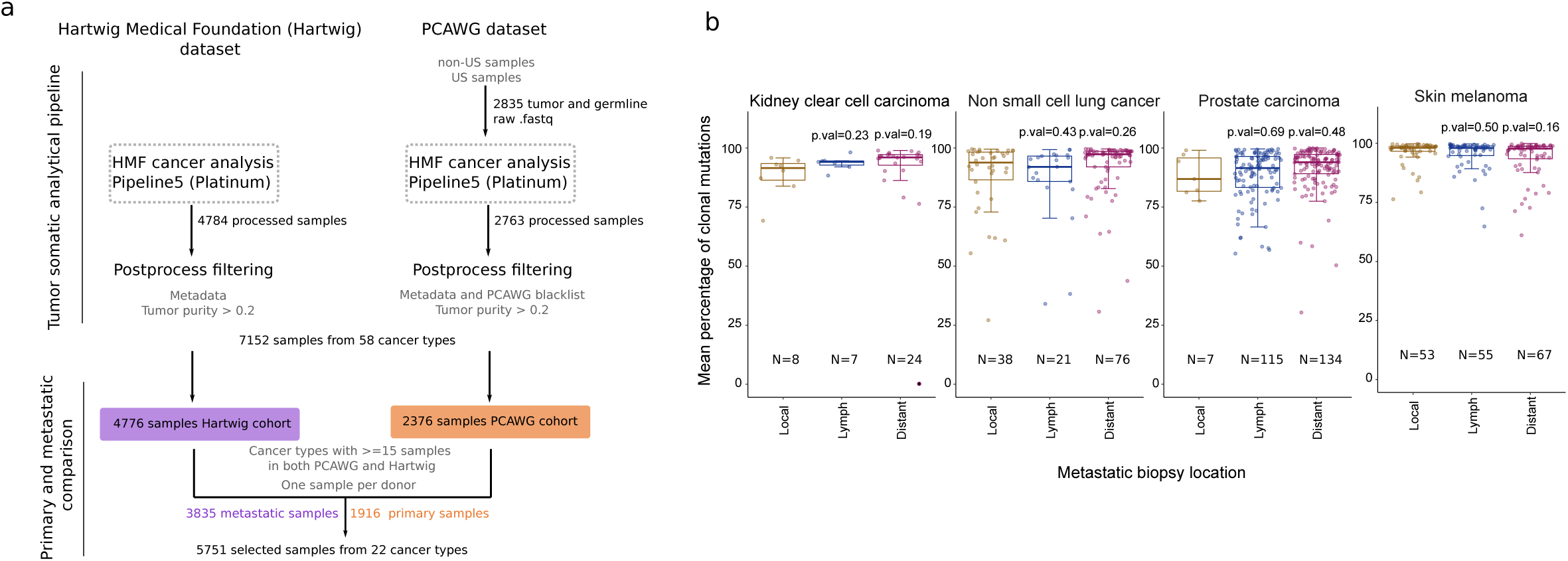
Cohort overview and global genomic features. **a)** Workflow of the unified processing pipeline used in this study for Hartwig (left) and PCAWG (right) WGS samples. First, PCAWG tumor and matched normal raw sequencing files were gathered and re-processed using the Hartwig tumor analytical pipeline. Next, the output of tumor samples that were correctly processed by the pipeline were further subjected to a strict quality control filtering. As a result, a total of 7,152 samples from 58 cancer types compose the harmonized dataset. 5,751 patient tumor samples from 22 cancer types with sufficient representation in both primary and metastatic datasets were selected for this study. **b)** Tumor clonality according to the metastatic biopsy location in kidney clear cell carcinoma, non-small cell lung cancer, prostate carcinomas and skin melanoma. N, number of samples in the group. p, Mann-Whitney p-value if p-value < 0.05. ns, not significant. Box-plots: center line, median; box limits, first and third quartiles; whiskers, lowest/highest data points at first quartile minus/plus 1.5× IQR.

**Supplementary Figure 2.**
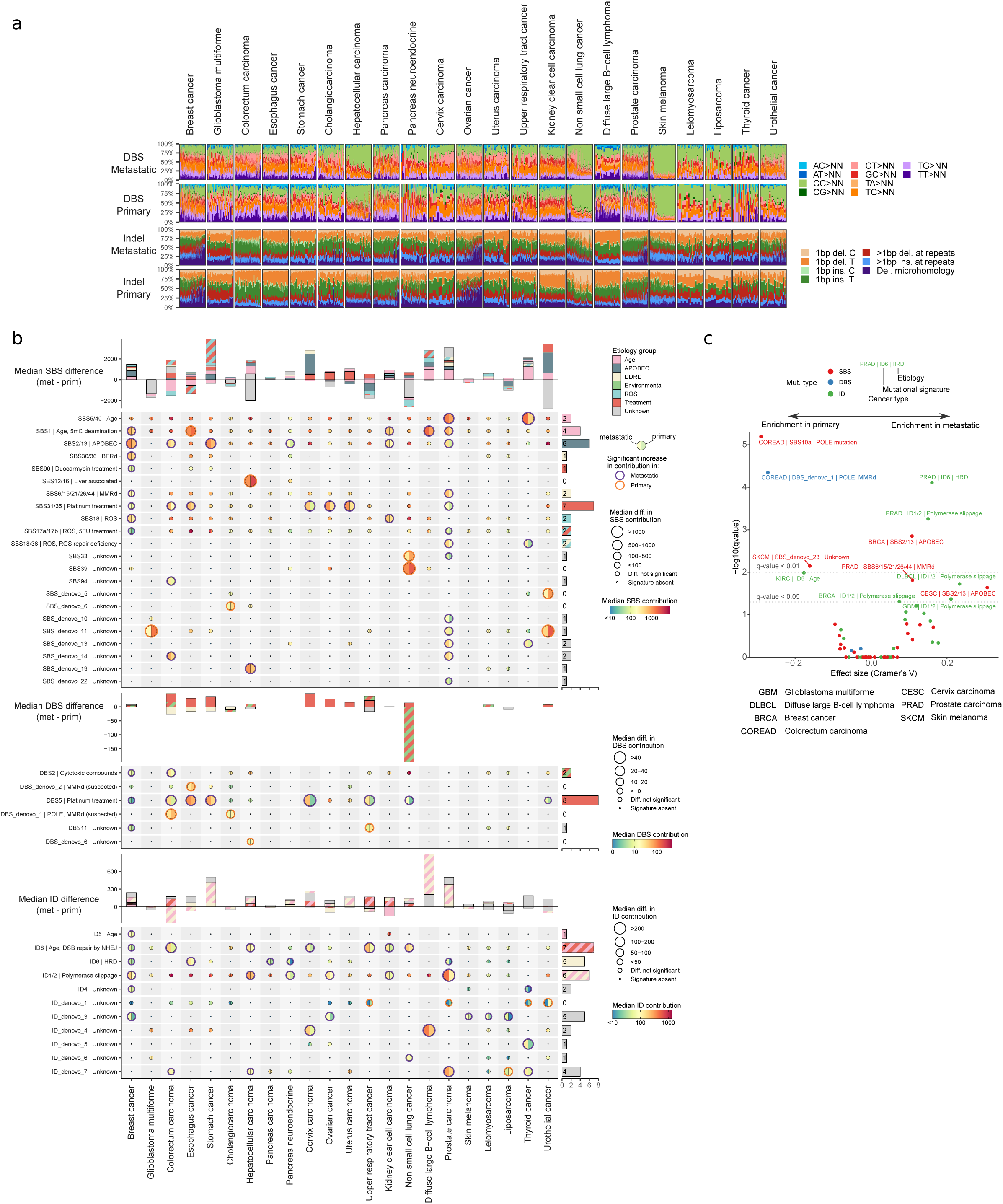
Mutation burden and mutational signatures. **a)** DBS (top) and ID (bottom) mutational spectra of metastatic and primary tumors. Patients are ordered according to their TMB burden. **b)** From top to bottom, moon plot representing the SBS, DBS and ID burden differences attributed to each mutational signature. Top stacked bars represent the cumulative signature exposure difference, including mutational signatures enriched in primary tumors (negative values). Thicker bar edge lines represent significance. Bars are coloured according to the annotated etiology. All mutational signatures, independent of their annotated etiology, are included. Diff., difference. Muts. mutations. Sig., mutational signature. Mut. mutational. Susp., suspected. **c)** Volcano plot representing the mutational signature hypermutation (>10,000 mutations for SBS, >500 for DBS, and >1000 for ID) prevalence comparison between primary and metastatic patients. Y-axis, log_10_(adjusted p-value). X-axis, effect size as Cramer’s V. Each dot represents a mutational signature in a cancer type. Dots are coloured according to the mutation type.

**Supplementary Figure 3.**
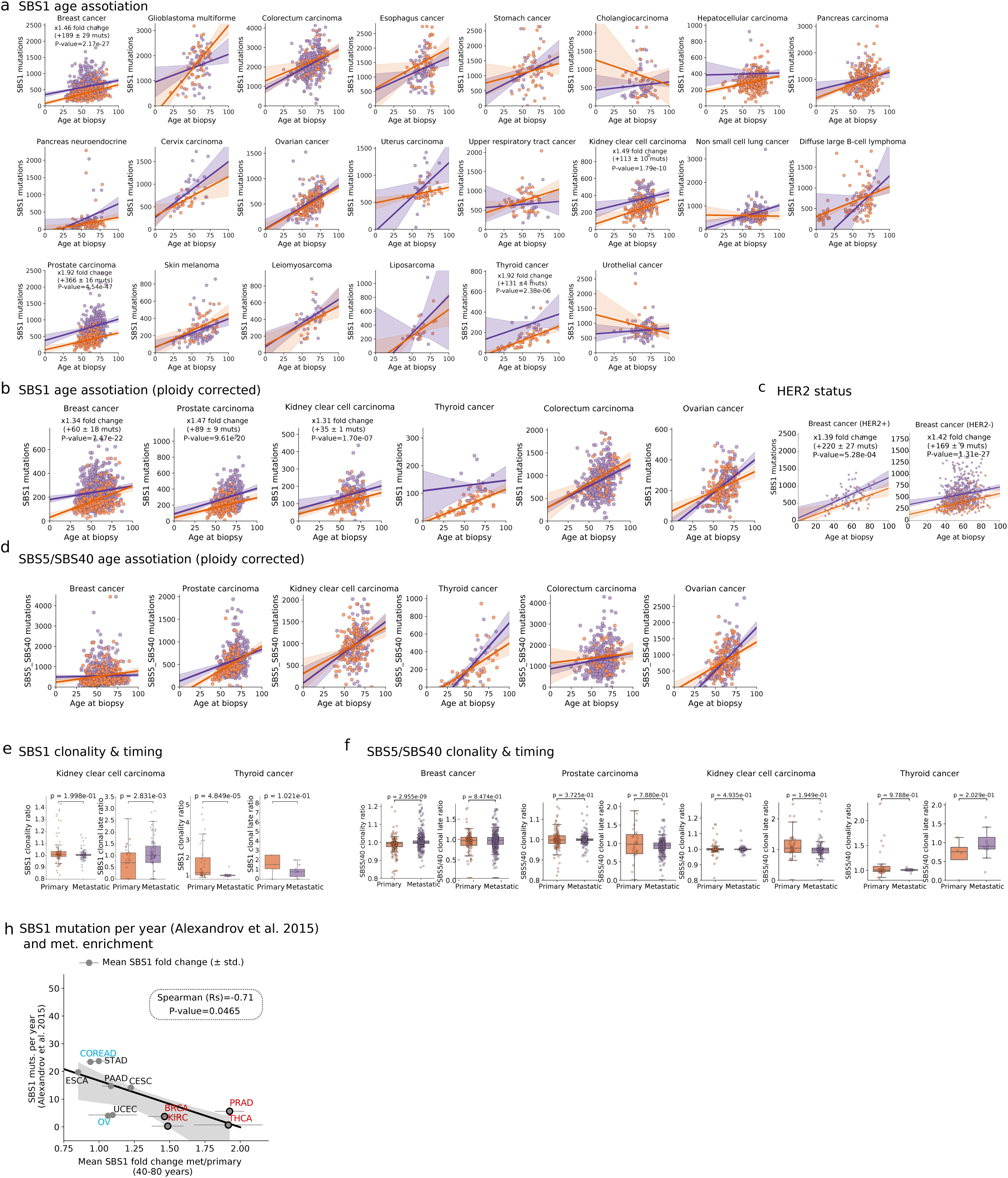
SBS1 mutation rates in primary and metastatic tumors. **a)** Linear regression of the SBS1 mutation burden (y-axis) and patient’s age at biopsy (x-axis) in primary and metastatic cancer across the 22 cancer types. The mean fold change, mean SBS1 increase per year and p-value are only represented in cancer types with an age-independent significantly different primary and metastatic distribution. **b)** Relative to a) for ploidy corrected SBS1 in the tumor types of interest. **c)** Independent linear regressions for HER2+ (left) and HER2 (right) breast cancer samples. ERBB2 amplification status was used to annotate cancer subtypes. **d)** Relative to a) for ploidy corrected SBS5/40 counts in the tumor types of interest. **e)** Equivalent to Fig. 3d for primary and metastatic in kidney clear carcinoma (left) and thyroid cancers (right). **f)** Relative to Fig. 3d, but using ploidy corrected SBS5/40 clonality ratio and clonal late ratio in the cancer types of interest. Boxplots are defined as in Fig. 1. **h)** Relative to Fig. 3f but using SBS1 year mutation rate from ref^45^. Muts, mutations.

**Supplementary Figure 4.**
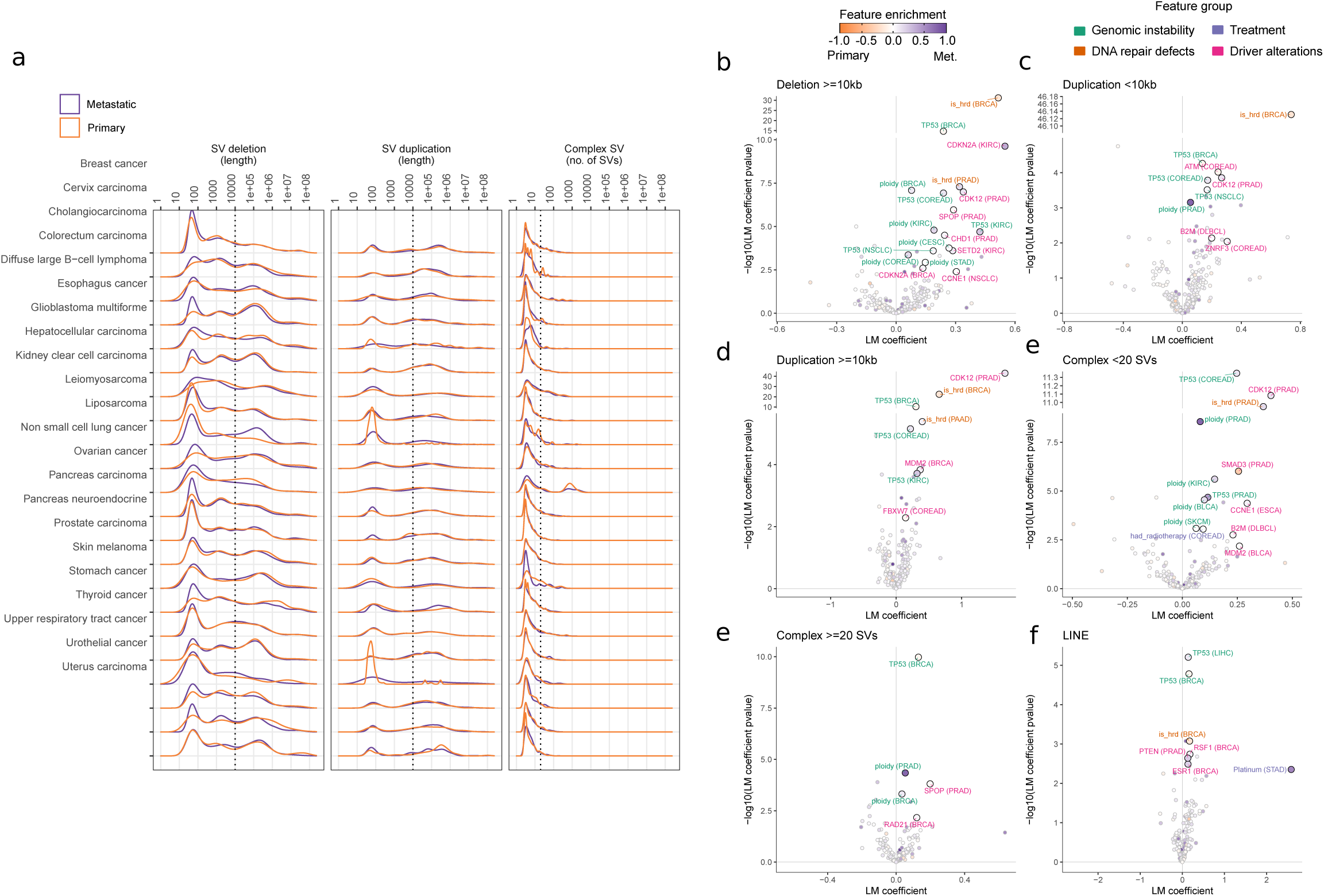
Structural variant burden. **a)** SV length frequency distribution of deletions (left panel) and duplications (middle panel). Right panel shows the frequency distribution of the number of linked breakpoints for complex SVs. Dashed vertical lines represent the chosen threshold to separate between short and large deletions, duplications and complex SVs, respectively. **b)** Volcano plot representing the cancer type specific regression coefficients (x-axis) and significance (y-axis) of clinical and genomic features against the number of large deletions. Each dot represents one feature in one cancer type. Labels are coloured according to the feature category. Dots are coloured by the frequency enrichment in metastatic (purple) or primary (orange) patients. LM, linear model. Coef, coefficient. Similar panels are displayed for **c)** short duplications, **d)** large duplications, **e)** short complex SVs, large complex SVs and **f)** LINEs.

**Supplementary Figure 5.**
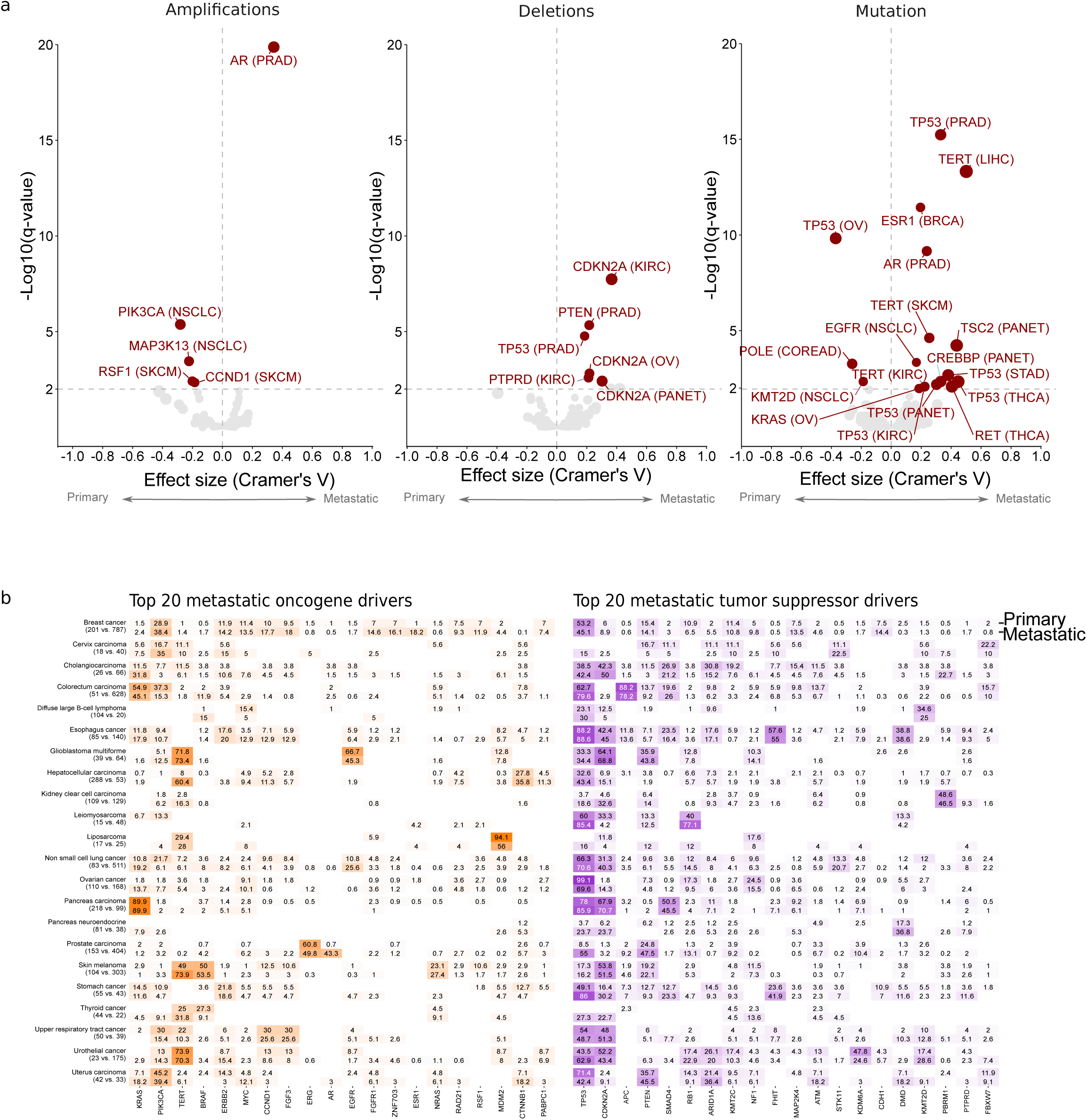
Driver landscape and drivers per patient. **a)** Volcano plots representing the cancer type specific enrichment (x-axis) and significance (y-axis) of driver genes between primary and metastatic cohorts. From left to right, amplification drivers, biallelically deleted drivers and mutated driver genes. BRCA, Breast cancer. CESC, Cervix carcinoma. CHOL, Cholangiocarcinoma. COREAD, Colorectal carcinoma. ESCA, Esophagus cancer. GBM, Glioblastoma multiforme. KIRC, Kidney clear cell carcinoma. LMS, Leiomyosarcoma. NSCLC, Non small cell lung cancer. OV, Ovarian cancer. PAAD, Pancreas carcinoma. PRAD, Prostate carcinoma. SKCM, Skin melanoma. STAD, Stomach cancer. THCA, Thyroid cancer. HNSC, Upper respiratory tract cancer. BLCA, Urothelial cancer. UCEC, Uterus carcinoma. **b)** Comparison fraction of mutated patients for the top 20 most frequently mutated (including all types of alterations) oncogenes (left) and tumor suppressor genes (right) in the metastatic cohort. Top numbers represent primary frequency, bottom metastatic.

**Supplementary Figure 6.**
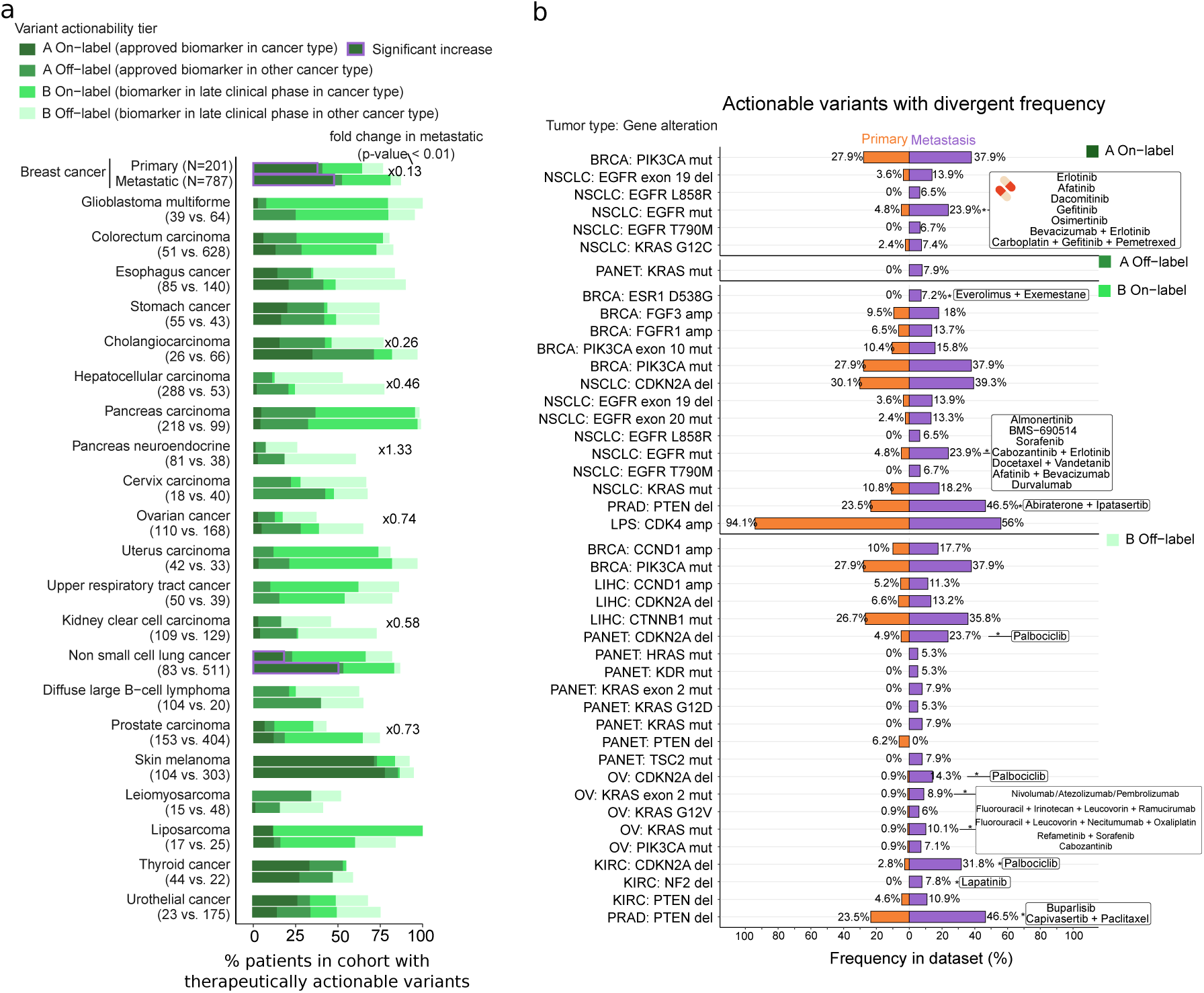
Therapeutic actionability of variants. **a)** Cancer type specific fraction of primary (top) and metastatic (bottom) patients with reported therapeutically actionable variants. For each patient the variant with the greatest level of evidence was considered. Bars are coloured according to the variant actionability tiers. Fold change labels are displayed in cancer types with a significant proportional increase (Fisher’s exact test p-value <0.01). Purple edgelines highlight significant increase in metastatic A-on label fraction patients. **b)** Primary (left) and metastatic (right) alteration frequency of actionable variants with a high discrepancy (>5% frequency difference) from cancer types with a global significant increase of actionable variants in metastatic patients from panel a). Asterisk, Fisher’s exact test p-value <0.01. Text boxes include the associated treatments for alterations with a significant mutation frequency increase in metastatic patients.

**Supplementary Figure 7.**
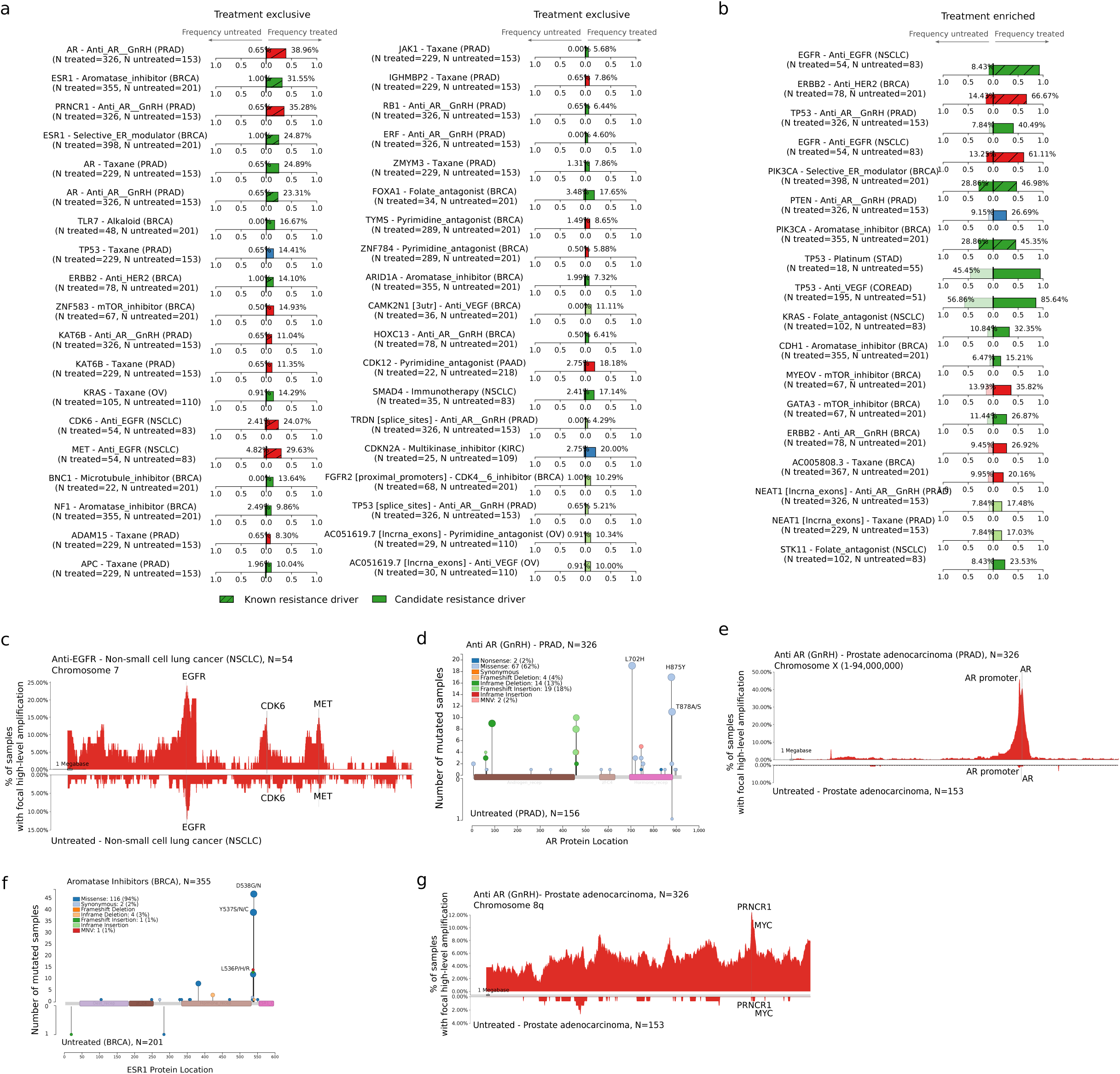
Treatment enriched drivers. **a)** Side by side alteration frequency comparison between treated (right bar) and untreated (left bar) patients for all treatment-exclusive and **b)** treatment enriched TEDs. **c)** Distribution of highly-focal copy number gains in chromosome 7 in non-small cell lung cancer untreated patients (bottom) and treated with anti-EGFR (top). *EGFR*, *MET* and *CDK6* genomic locations are highlighted. Each bin represents 100Kbs. **d)** Distribution of mutations along the *AR* protein sequence in prostate cancer patients treated with androgen deprivation (top) and untreated (bottom). Pfam domains are represented as rectangles. Mutations are coloured according to the consequence type. **e)** Distribution of highly-focal copy number gains in chromosome X in prostate untreated patients (bottom) and treated with androgen deprivation(top). *AR* coding region and the promoter region are highlighted. Each bin represents 100Kbs. **f)** Distribution of mutations along the *ESR1* protein sequence in breast cancer patients treated with aromatase inhibitors (top) and untreated (bottom). **g)** Similar to e) but representing MYC and PRNCR1 co-amplifications in chromosome 8q.

### Supplementary Tables

**Supplementary Table 1.** Primary and metastatic cohorts metadata.

**Supplementary Table 2.** Karyotype and genomic instability measurements.

**Supplementary Table 3.** Mutational signatures.

**Supplementary Table 4.** SBS1 mutation rate.

**Supplementary Table 5.** Structural variants.

**Supplementary Table 6.** Driver alterations.

**Supplementary Table 7.** Therapeutic actionability of variants.

**Supplementary Table 8.** Treatment associated drivers (TEDs).

### Supplementary Notes

**Supplementary Note 1.** Application of Hartwig analytical pipeline to the PCAWG dataset.

### Methods

#### Cohort gathering and processing

We have matched tumor-normal whole genome sequencing data from cancer patients from two cohorts: the Hartwig Medical Foundation (Hartwig) and the Pan-Cancer Analysis of Whole Genomes (PCAWG) cohort. A detailed description of the Hartwig and PCAWG cohort gathering and processing as well as comprehensive documentation of the PCAWG sample reanalysis with the Hartwig somatic pipeline is described in the Supp. Note 1.

#### Tumor clonality analysis

Each mutation in the .vcf files is given a subclonal likelihood by PURPLE. Following PURPLE guidelines, we considered mutations with subclonal scores of 0.8 or higher to be subclonal and mutations below the 0.8 threshold to be clonal. For each sample we then computed the average proportion of clonal mutations by dividing the number of clonal mutations by the total mutation burden (including SBS, MNVs and indels). Finally, for each cancer type we used Mann-Whitney test to assess the significance of the clonality difference between the primary and metastatic tumors. A p-value lower than 0.05 was deemed as significant.

In addition, we leveraged biopsy site data in patient reports to further investigate differences in metastatic tumor clonality according to the metastatic biopsy site. If the metastatic biopsy site was in the same organ/tissue as the primary tumor, we considered them as “Local”, while if the metastatic biopsy site was reported in the lymphoid system or other organs/tissues they were dubbed as “Lymph” and “Distant”, respectively. Cancer types for which there was a minimum of 5 samples available for each of the biopsy groups were selected and Mann-Whitney test was used to compare the clonality between the biopsy groups.

#### Karyotype

Arm-level and genome copy number (CN) was estimated as described in Taylor et al. 2018^31^. Briefly, CN of segments determined by PURPLE were rounded to the nearest integer, then the arm coverage per CN was calculated. Arm-level CN was determined to be either the CN with coverage >50%, the most common arm CN if coverage <50%, or the genome CN if coverage <50% and the most common arm CN does match the genome CN. The most common arm CN across all arms was deemed to be the genome ploidy.

Estimated genome ploidy per sample was subtracted from the estimated chromosome arm ploidy to create a matrix of chromosome arm gains and losses relative to the estimated genome ploidy. This matrix was stratified by cohort (metastatic/primary) and a Mann-Whitney test was performed to assess the difference of the distributions for each arm in each cancer type. The resulting p-value was FDR adjusted across all arms per cancer type. A q-value < 0.05 was deemed to be significant. Mean arm gains and losses relative to genome ploidy were calculated and represented.

#### Genomic instability markers

To compare the differences in aneuploidy scores and the LOH proportions in each group, a Mann-Whitney test was performed per cancer type. The aneuploidy score represents the number of arms per tumor sample that deviate from the estimated genome ploidy as described in Taylor et al. 2018^31^. To compare the LOH proportions, only diploid samples were included when calculating the p-value. The LOH proportion was defined as 1 - (diploid proportion estimated by PURPLE) of the genome.

To compare the fraction of samples with a driver mutation in TP53 as well as the fraction of whole genome duplicated samples per cohort, a Fisher’s exact test was performed per cancer type. Any TP53 driver alteration (non-synonymous mutation, biallelic deletion and homozygous disruption) was considered in the analysis. Whole genome duplication was defined as present if the sample had more than 10 autosomes with an estimated chromosome copy number >1.5. A p-value < 0.01 was deemed to be significant for all statistical tests.

#### Mutational signature analysis

##### Signature extraction

The number of somatic mutations falling into the 96 single nucleotide substitution (SBS), 78 double base substitutions (DBS) and 83 indel (ID) contexts (as described in the COSMIC catalog^75^ https://cancer.sanger.ac.uk/signatures/) was determined using the R package mutSigExtractor (https://github.com/UMCUGenetics/mutSigExtractor, v1.23).

SigProfilerExtractor (v1.1.1) was then used (with default settings) to extract a maximum of 21 SBS, 8 DBS and 10 ID *de-novo* mutational signatures. This was performed separately for each of the 22 tissue types which had at least 30 patients in the entire dataset (aggregating primary and metastatic samples, see Supp. Table 3). Tissue types with less than 30 patients as well as metastatic patients with unknown primary location type were combined into an additional ‘Other’ group, resulting in a total of 23 tissue type groups for signature extraction. In order to select the optimum rank (i.e. the eventual number of signatures) for each tissue type and mutation type, we manually inspected the average stability and mean sample cosine similarity plots output by SigProfilerExtractor. This resulted in 440 *de-novo* signature profiles extracted across the 23 tissue type groups (Supp. Table 3). Least squares fitting was then performed (using the fitToSignatures() function from mutSigExtractor) to determine the per-sample contributions to each tissue type specific *de-novo* signature.

##### Etiology assignment

The extracted *de-novo* mutational signatures with high cosine similarity (≥0.85) to any reference COSMIC mutational signatures with known cancer type associations^75^ were labeled accordingly (256 *de-novo* signatures matched to 56 COSMIC reference signatures).

For the remaining 184 unlabeled *de novo* signatures, we reasoned that there could be one or more signatures from one cancer type that are highly similar to those found in other tissue types, and that these likely represent the same underlying mutational process. We therefore performed clustering to group likely equivalent signatures. Specifically, the following steps were performed:

1. We calculated the pairwise cosine distance between each of the *de-novo* signature profiles.
2. We performed hierarchical clustering and used the base R function cutree() to group signature profiles over the range of all possible cluster sizes (min no. clusters = 2; max no. of clusters = number of signature profiles for the respective mutation type).
3. We calculated the silhouette score at each cluster size to determine the optimum number of clusters.
4. Finally, we grouped the signature profiles according to the optimum number of clusters. This yielded 27 SBS, 7 DBS, and 8 ID *de-novo* signature clusters (see Supp. Table 3).

For certain *de novo* signature clusters, we could manually assign the potential etiology based on their resemblance to signatures with known etiology described in COSMIC^75^, Kucab *et al.* 2019^41^ and Signal ^76^. Some clusters were an aggregate of 2 known signatures, such as SBS_denovo_clust_2 which was a combination of SBS2 and SBS13, both linked to APOBEC mutagenesis. Other clusters had characteristic peaks of known signatures, such as DBS_denovo_clust_4 which resembled DBS5 based having distinct CT>AA and CT>AC peaks. Lastly, DBS_denovo_clust_1 was annotated as POLE mutation and MMR deficiency as samples with high contribution (>150 mutations) of this cluster are frequently MSI or have POLE mutations. Likewise, DBS_denovo_clust_2 was annotated with MMR deficiency as the etiology as samples with high contribution (>250 mutations) of this cluster were all MSI. See Supp. Table 3 for a list of all the manually assigned etiologies.

##### Comparing the prevalence of mutational processes between primary and metastatic cancer

We then compared the activity (i.e. number of mutations contributing to) of each mutational process between primary and metastatic tumors. For each sample, we first summed the contributions of signatures of the same mutation type (i.e. SBS, DBS or ID) with the same etiology, henceforth referred to as ‘etiology contribution’. Per cancer type and per etiology, we performed two-sided Mann-Whitney tests to determine whether there was a significant difference in etiology contribution of primary and metastatic tumors. Per cancer type and per mutation type, we used the p.adjust() base R function to perform multiple testing correction using Holm’s method. Next, we added a pseudocount of 1 to the contributions (to avoid dividing by zero) and calculated the median contribution log2 fold change, i.e. log2[ (median contribution in metastatic tumors + 1) / (median contribution in primary tumors + 1) ]. We considered the etiology contribution between primary and metastatic tumors to be significantly different when the q-value was <0.05, and log_2_ fold change was ≥0.4 or ≤ -0.4 (= ±x1.4).

We also determined whether there was an increase in the number of samples with high etiology contribution (i.e., hypermutators) in metastatic versus primary cohorts. For each signature, a sample was considered a hypermutator if the etiology contribution was ≥10,000 for SBS signatures, ≥500 for DBS signatures, or ≥1,000 for ID signatures. For each cancer type, for each etiology, we performed pairwise testing only for cases where there were ≥5 hypermutator samples for either metastatic or primary tumors. Each pairwise test involved calculating p-values using two-sided Fisher’s exact tests, and effect sizes by multiplying Cramer’s V by the sign of the log2(odds ratio) to calculate a signed Cramer’s V value that ranges from -1 to +1 (indicating enrichment in primary or metastatic respectively). We then used the p.adjust() base R function to perform multiple testing correction using Bonferroni’s method.

#### SBS1-age correlations in primary and metastatic tumors

To count the SBS1 mutations we relied on the definition from ref^77^ that is based on the characteristic peaks of COSMIC SBS1 signature profile: single base CpG > TpG mutations in NpCpG context. To ensure that these counts and the downstream analyses are not affected by differential APOBEC exposure in primary and metastatic cohorts, we excluded CpG > TpG in TpCpG which is also a characteristic peak in COSMIC SBS2 signature profile. Also, for skin melanoma CpG > TpG in [C/T]pCpG which overlaps with SBS7a was excluded. To obtain the SBS5 and SBS40 counts we relied on their exposures derived from the mutational signature analyses performed in this study (explained above).

To assess the correlation between SBS1 burden and patient’s age at biopsy we performed a cancer type and cohort specific linear regression (i.e., separate regression for primary and metastatic tumor samples). To avoid spurious effects caused by hypermutated tumors, samples with TMB greater than 30,000 as well as those with SBS1 burden greater than 5,000 were excluded.

For each cancer type and cohort we then computed 100 independent linear regressions by randomly selecting 75% of the available samples. We selected the median linear regression (based on the regression slope) as representative regression for further analyses. Similarly, confidence intervals were derived from the 1st and 99th percentile of the computed regressions.

To evaluate the significance of the differences between primary and metastatic representative linear regressions (hereafter referred to as linear regression for simplicity) we first filtered out cancer types that failed to show a positive correlation trend between SBS1 burden and age at biopsy in both primary and metastatic tumors (i.e., Pearson’s correlation coefficient of primary and metastatic regression >0.1). Next, for each selected cancer type, we computed the regression residuals of primary and metastatic SBS1 mutation counts using, in both cases, the primary linear regression as baseline. The primary and metastatic residual distributions were then compared using a Mann-Whitney test to evaluate significance. Cancer types with a Mann-Whitney p-value < 0.01 were deemed as significant. Finally, to ensure that the differences were uniform across different age ranges (i.e., not driven by a small subset of patients) we only considered significant cancer types where the metastatic linear regression intercept is higher than the primary intercept.

SBS5/SBS40 correlations were computed following the same procedure and using the sum of SBS5 and SBS40 exposures for each tumor sample. If none of the mutations were attributed to SBS5/SBS40 mutational signatures, the aggregated value was set to zero. In the ploidy corrected analyses we divided the SBS1 mutation counts (and SBS5/SBS40 mutation counts for the SBS5/40 ploidy corrected regression, respectively) by the PURPLE estimated tumor genome ploidy.

For each cancer type the mean fold-change (fc) was defined as 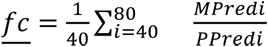 where MPred_i_ and PPred_i_ are the estimated number of SBS1 mutations for a given age i-th according to the metastatic and primary linear regressions, respectively. Similarly, the mean estimated SBS1 burden difference (SBS1_diff_) was defined as: 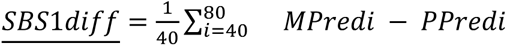.

#### Clonality and timing of clock-like mutations

SBS1 individual mutations were identified as described in the previous section. For SBS5 and SBS40 mutations, we used a maximum likelihood approach to assign individual mutations to the SBS5 and SBS40 mutational signatures in a cancer type specific manner.

For every SBS1 (and SBS5/SBS40 mutation) we then assign the clonality according to the PURPLE subclonal likelihood estimation, where only mutations with SUBCL likelihood >= 0.8 were considered as such (see above). Likewise, the molecular timing of individual mutations (i.e., clonal early and clonal late) was computed using the MutationTimeR^77^ package.

For each tumor sample the SBS1 clonality ratio (or respectively SBS5/SBS40 clonality ratio) was defined as the ratio between the proportion of clonal SBS1 mutations 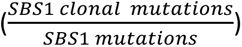 divided by the total proportion of clonal alterations in the sample 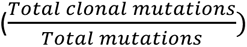. Similarly, the SBS1 clonal late ratio was defined as the ratio between the proportion of clonal late SBS1 mutations 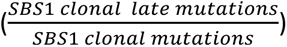 divided by the total proportion of clonal late alterations in the sample 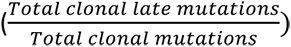, where the total of clonal SBS1/total mutations was computed as the sum of clonal late, clonal early and unassigned clonal from MutationTimerR.

#### Primary cell division rate and accelerated SBS1 mutagenesis in metastasis

To assess the relationship of cell division rate of primary tumors with the accelerated SBS1 mutagenesis in the metastatic setting we relied on the SBS1 burden per year as a proxy of stem-cell division rates as was previously described in ref^45^. We thus computed for each primary cancer type the average number of SBS1 per year as the number of SBS1 mutations divided by the patient’s age at biopsy (only considering primary samples and excluding hypermutated samples as described above). We then used a Spearman’s correlation to assess its association with the estimated mean SBS1 mutation rate fold change in metastatic tumors (see above). Additionally, to exclude potential biases in our primary cohort, we repeated the same analysis relying on an independent measurement of primary cancer SBS1 yearly accumulation. Specifically, we used the best estimated accumulation of SBS1 per year from ref^45^ Supplementary Data Set 6 and regressed it to the fold change estimates for the matching cancer types present in both datasets.

#### Structural variant (SV) analysis

##### SV type definitions

LINX^53^ chains one or more SVs and classifies these SV clusters into various event types (‘ResolvedType’). We defined deletions and duplications as clusters with a ResolvedType of ‘DEL’ or ‘DUP’ whose start and end breakpoints are on the same chromosome (i.e. intrachromosomal). Deletions and duplications were split into those <10kb and ≥10kb in length (small and large, respectively), based on observing bimodal distributions in these lengths across cancer types (Supp. Fig. 4a). We defined complex SVs as clusters with a ‘COMPLEX’ ResolvedType, an inversion ResolvedType (including: INV, FB_INV_PAIR, RECIP_INV, RECIP_INV_DEL_DUP, RECIP_INV_DUPS), or a translocation ResolvedType (including: RECIP_TRANS, RECIP_TRANS_DEL_DUP, RECIP_TRANS_DUPS, UNBAL_TRANS, UNBAL_TRANS_TI). Complex SVs were split into those with <20 and ≥20 SVs (small and large, respectively), based on observing similar unimodal distributions in the number SVs across cancer types whose tail begins at ∼20 breakpoints (Supp. Fig. 4a). Lastly, we defined LINEs (long interspersed nuclear element insertions) as clusters with a ResolvedType of ‘LINE’. For each sample, we counted the occurrence (i.e. SV burden) of each of the 7 SV types described above. Additionally, we determined the total SV burden by summing counts of the SV types.

##### Comparing SV burden between primary vs. metastatic cancer

We then compared the SV type burden between primary versus metastatic tumors as shown in Fig. 4a. Firstly, we performed Mann-Whitney tests per SV type and per cancer type to determine whether there was a significant difference in SV type burden between primary versus metastatic, and used p<0.01 as the significance threshold. Next, we calculated relative enrichment as follows: log_10_(median SV type burden in metastatic tumors + 1) − log_10_(median SV type burden in primary tumors + 1); and calculated fold change as follows: (median SV type burden in metastatic tumors + 1) / (median SV type burden in primary tumors + 1). When calculating relative enrichment and fold change, the pseudocount of 1 was added to avoid the log(0) and divide by zero errors, respectively. Fold changes are displayed with a ‘>’ in Fig. 4a when the SV burden for primary tumors is 0 (i.e. when a divide by zero would occur without the pseudocount).

##### Identifying features associated with SV burden increase in metastatic cancer

To identify the features that could explain increased SV burden, we correlated SV burden with various tumor genomic features. This included: i) genome ploidy (determined by PURPLE); ii) HRd status (determined by CHORD^50^) and MSI status (determined by PURPLE); iii) the presence of mutations in 345 cancer associated genes (excluding fragile site genes that are often affected by CNVs^14^), henceforth referred to as ‘gene status’; and iv) treatment history, including the presence of radiotherapy, the presence of one of the 79 different cancer therapies as well as the total number of treatments received. All primary samples as well as all metastatic samples without treatment information were considered to have no treatment. Genome ploidy and total number of treatments received were numeric features, whereas all of the remaining were boolean (i.e. true/false) features. In total there were 429 features.

SV type burden was transformed to log_10_(SV type burden + 1) and was correlated with the 429 features using multivariate linear regression models (LM). This was performed separately for each of the 22 cancer types, and for each of the 7 SV types, resulting in a total of 154 (=22 cancer types x 7 SV types) LM models.

Each LM model (i.e. per SV type and cancer type) involved training of three independent LMs with i) both metastatic and primary samples (primary+metastatic), ii) only Hartwig samples (metastatic-only), and iii) only primary samples (primary-only). This was done to filter out correlations between features and increased SV type burden solely due to differences in feature values between primary and metastatic tumors. We then required features that positively correlated with SV type burden in the primary+metastatic LM to independently show the same association in the metastatic-only or primary-only LMs. Only genomic features that independently showed positive correlation with the SV burden were further considered as significant (i.e., represented in the lollipots).

Each of the 3 LMs was trained as follows:

1. Remove boolean features with too few ‘true’ samples:

a. For the primary+metastatic LM, remove gene status features with <15 ‘true’ samples.
b. For the metastatic-only and primary-only LMs, remove gene status features with <10 ‘true’ samples.
c. For the remaining boolean features, remove features with <5% ‘true’ samples.
2. Fit a LM using the lm() base R function to correlate log_10_(SV type burden + 1) versus all features.

For each LM analysis, we used the following filtering criteria to identify the features that were correlated with increased SV type burden:

1. Only keep LM analyses where there was significant increase in SV type burden for the respective cancer type (p<0.01 as described in the previous section “Comparing SV burden between primary vs. metastatic cancer”)
2. For primary+metastatic LM

a. Regression p-value <0.01
b. Coefficient p-value <0.01
c. Coefficient >0
3. For metastatic-only LM or primary-only LM:

a. Coefficient p-value <0.01
b. Coefficient >0

Finally, to determine which features (of those correlated with increased SV type burden) were enriched in metastatic tumors compared to primary tumors (and vice versa), we calculated Cliff’s delta for numeric features and Cramer’s V for boolean features. Cliff’s delta ranges from -1 to +1, with -1 representing complete enrichment in primary tumors, whereas +1 represents complete enrichment in metastatic tumors. Cramer’s V only ranges from 0 to 1 (with 1 representing enrichment in either primary or metastatic tumors), the sign of the log(odds ratio) was assigned as the sign of the Cramer’s V value so that it ranges from -1 to +1. Features with an effect size >0 were considered as those that could explain the SV burden increase in metastatic cancer when compared to primary cancer.

#### Driver alterations

We relied on patient specific cancer driver catalogs constructed by PURPLE^14^ and LINX^53^. Only drivers with a driver likelihood > 0.5 were retained. Fusion drivers were filtered for those that were previously reported in the literature. Similarly, we manually curated the list of drivers and removed SMAD3 HOTSPOT mutations because of the high burden mutations in low mappability regions. The final driver catalog contained a total of 443 driver genes.

We then combined fusions with the LINX driver variants to calculate a patient specific number of driver events. Drivers that concern the same driver gene but a different driver type were deemed to be individual drivers (e.g. TP53 mutation and TP53 deletion in the same sample were considered as one driver event). Cancer type specific Mann-Whitney test was performed to assess differences between primary and metastatic tumors. A p-value < 0.01 was deemed to be significant.

A contingency matrix was constructed from the driver catalog, containing the frequency of driver mutations per driver type (i.e., deletion, amplification or mutations) and cancer type in each cohort (metastatic and primary). A second contingency matrix was constructed for the fusions. Partial amplifications were considered as amplifications while homologous disruptions were considered as deletions. These contingency matrices were filtered for genes which show a minimum frequency of 4 mutated samples in either the primary or the metastatic cohorts. Then a Fisher’s exact test for each gene, cancer type and mutation type was performed and the p-value was adjusted for FDR per cancer type. Cramer’s V and the odds ratio were used as effect size measures. An adjusted p-value < 0.01 was deemed to be significant.

#### Therapeutic actionability of variants

To determine the amount of actionable variants observed in each sample, we compared our variants annotated by SnpEff^78^ to those derived from three different databases (OncoKB^79^, CIViC^80^ and CGI^81^) that were classified based on a common clinical evidence level (https://civic.readthedocs.io/en/latest/model/evidence/level.html) as described in Priestley et al. 2019^14^. In our study we only considered A and B levels of evidence which represent variants that have been FDA approved for treatment and are currently being evaluated in a late stage clinical trial, respectively. A variant was determined to be “On-label” when the cancer type it was found in matches the cancer type the treatment was approved for or is being investigated for, and “Off-label” otherwise. Only actionable variants of the sensitive category were considered (i.e. tumors containing the variant are sensitive to a certain treatment). Sample-level actionable variants such as TMB high/low or MSI status were not evaluated, because of their tendency to overshadow the other variants, especially in the Off-label category. Further, wild-type actionable variants were not considered in this analysis for the same reason. Variants related to gene expression or methylation were not considered due to lack of available data. Additionally, we found actionable variants derived from leukemias to be very different from the solid tumors in our data set which is why we excluded them for this analysis. For the analysis of proportion of samples bearing therapeutically actionable variants we considered the highest evidence level was retained for each sample following the order A On/Off-label to B On/Off-label. To assess enrichment of actionable variants globally and on A On-label level in metastatic tumors, a Fisher’s exact test was performed pancancer-wide and per cancer type. A p-value < 0.01 was deemed to be significant. Percentage changes in frequency are only shown for significant cases.

To determine which variants contribute the most to the observed significant frequency differences, individual actionable variants were tested for enrichment in metastatic tumors using a Fisher’s exact test per cancer type and tier level. P-values were FDR adjusted per cancer type and a q-value < 0.05 was deemed to be significant. In Supp. Fig. 6 only actionable variants that were found at a minimum frequency of 5% in either primary or metastatic cohort and a minimum frequency difference of 5% between them were shown. However, the differences across all screened variants is available as part of Supp. Table 7.

#### Treatment enriched drivers

We aimed to pinpoint drivers that are potentially responsible for lack of response to certain cancer treatments in the metastatic cohort. Hence, we devised a test that identifies driver alterations that are enriched in groups of patients treated with a particular treatment type compared to the untreated group of patients from the same cancer type (see Fig. 6a for illustration of the workflow).

Treatments were grouped according to their mechanism of action so that multiple drugs with a shared mechanism of action were grouped into the mechanistic treatment category (e.g. Cisplatin, Oxaliplatin, Carboplatin as Platinum). We created 332 treatment and cancer type groups by grouping patients with treatment annotation according to their treatment record before the biopsy. One patient might be involved in multiple groups if they have received multiple lines of therapy or a simultaneous combination of multiple drugs. Only 62 treatment and cancer type groups with at least 10 patients were further considered in the analysis.

Hence, for each cancer and treatment group we performed the following steps:

1. We first performed a driver discovery analysis in treatment and cancer type specific manner. We explored three types of somatic alterations: coding mutations, non-coding mutations and copy number variants (see below for detailed description of each driver category). Driver elements from each alteration category were selected for further analysis.
2. For each driver alteration from 1) we compared the alteration frequency in the treated group to the untreated group of the same cancer type. Each driver category (coding and non coding mutations and copy number variants) were evaluated independently. We performed a Fisher’s exact test to assess the significance of the frequency differences. Similarly, we computed the odds ratio of the mutation frequencies for each driver alteration. The p-values were adjusted with a multiple-testing correction using the Benjamini–Hochberg procedure (alpha=0.05). An adjusted p-value of 0.05 was used for coding mutations and copy number variants. Adjusted p-value of 0.1 was used for non-coding variants due to the overall low mutation frequency of the elements included in this category, which hampered the identification of significant differences.
3. We then annotated each driver element with information about the exclusivity in the treatment group. We labeled drivers as treatment exclusive if the mutation frequency in the untreated group was lower than 5%, we annotated as treatment enriched otherwise. Additionally, we manually curated the identified drivers with literature references of their association with each treatment category.
4. Finally, the overlap of patients in multiple treatment groups (see above) in the same cancer type prompted us to prioritize the most significant treatment association for each driver gene in a particular cancer type. In other words, for each driver gene that was deemed as significantly associated with multiple treatment groups in the same cancer type, we selected the most significant treatment association, unless a driver-treatment annotation was clearly reported in the literature.

The full catalog of TEDs and their mutation frequencies can be found in Supp. Table 8.

##### Coding mutations drivers

We used dNdScv^82^ with default parameters to identify cancer driver genes from coding mutations. A global q-value <0.1 was used as a threshold for significance. Mutation frequencies for each driver gene were extracted from the dndscv output. We defined the mutation frequency as the number of samples bearing non-synonymous mutations.

##### Non-coding mutations drivers

We used ActiveDriverWGS^83^ (v1.1.2, default parameters) to identify non-coding driver elements in five regulatory regions of the genome including 3’UTRs, 5’UTRs, lncRNAs, proximal promoters and splice sites. For each element category we extracted the genomic coordinates from Ensembl v101. Each regulatory region was independently tested. To select for significant hits, we filtered on adjusted p.values (FDR<0.1) and minimum of three mutated samples. We defined the mutation frequency as the number of mutated samples for each significantly mutated element in the treatment group.

##### Copy number variant drivers

We ran GISTIC2^84^ (v2.0.23) on each of the 62 treatment and cancer type groups using the following settings:

gistic2 -b <INPUTPATH> -seg <INPUTSEGMENTATION> -refgene hg19.UCSC.add_miR.140312.refgene.mat -genegistic 1 -gcm extreme -maxseg 4000 - broad 1 -brlen 0.98 -conf 0.95 -rx 0 -cap 3 -saveseg 0 -armpeel 1 -smallmem 0 -res 0.01 -ta 0.1 -td 0.1 -savedata 0 -savegene 1 -qvt 0.1.

The focal GISTIC peaks (*q* ≤ 0.1 and < 1Mbp) were then annotated with functional elements using the coordinates from Ensembl v101. The frequency differences between treated and untreated cohorts on every gene was assessed with Fisher exact test as described above. For this, we first calculated the focal amplification and deep depletion status of every gene within each sample. A gene was amplified when the ploidy level of the gene was 2.5 ploidy levels higher than its genome-wide mean ploidy level (as measured by PURPLE), and deleted when the gene ploidy level was lower than 0.3 (i.e. deep deletion). We observed that the majority of the peaks contained multiple significant gene candidates (after multiple correction q < 0.05) and therefore we retained the gene most closely positioned to the peak summit, which is the most significantly enriched region across the treated samples. Next, we also found recurrent peaks across multiple treatment groups per cancer type that are not, or less, present in the untreated control group because most of the Hartwig samples have received multiple treatment types. We therefore merged peaks with overlapping ranges to produce a single peak per genomic region per cancer type. For each collapsed peak we selected the treatment type showing the lowest q-value for the gene near the peak summit. Deletion and amplification peaks were processed separately.

#### Group level aggregation of treatment resistance associated variants

To estimate the contribution of TEDs to the total number of drivers per sample in the metastatic cohort, we excluded any TED from the catalog of driver mutations (see above Driver alterations section) in a cancer type, gene and driver type specific manner.

## Data availability

Metastatic WGS data and metadata from the Hartwig Medical Foundation are freely available for academic use through standardized procedures. Request forms can be found at https://www.hartwigmedicalfoundation.nl.

Somatic variant calls, gene driver lists, copy number profiles and other core data of the PCAWG cohort generated by the Hartwig analytical pipeline are available for download at https://dcc.icgc.org/releases/PCAWG/Hartwig. Researchers will need to apply to the ICGC data access compliance office (http://icgc.org/daco) for the ICGC portion of the dataset. Authentication of NIH eRA commons is required to access the TCGA portion of the dataset via https://icgc.bionimbus.org. Additional information on accessing the data, including raw read files, can be found at https://docs.icgc.org/pcawg/data/.

## Code availability

The Hartwig analytical processing pipeline is available at (https://github.com/hartwigmedical/pipeline5) and implemented in Platinum (https://github.com/hartwigmedical/platinum).

The source code to reproduce the analysis of the manuscript will be made public in this repository https://github.com/UMCUGenetics/PCAWG_Hartwig_comparison upon manuscript acceptance in a peer-review journal.

## Author contributions

Conceptualization, FMJ, AvH and EC. Methodology, FMJ, AM, SB, LN, PP, AvH. Software, FMJ, AM, SB, LN, PP, AvH. Validation, FMJ, AM, SB, LN, PP, AvH. Formal Analysis, FMJ, AM, SB, LN, PP, AvH and EC. Investigation Resources, FMJ, AM, SB, LN, PP, AvH . Data Curation, FMJ, AM, SB, LN, PP, AvH. Writing Original Draft, FMJ and AvH. Writing - Review & Editing, FMJ, AM, SB, LN, PP, AvH and EC. Visualization, FMJ, AM, SB, LN, AvH and EC. Supervision, FMJ, AvH and EC. Project Administration, AvH and EC. Funding Acquisition EC.

## Acknowledgements

This publication and the underlying study have been made possible partly on the basis of the data that Hartwig Medical Foundation and the Center of Personalised Cancer Treatment (CPCT) have made available to the study.

We thank Roel Janssen for the technical assistance in the collection and processing of the PCAWG raw sequencing data using the Hartwig tumor analytical pipeline. We thank Joaquin Mateo for his valuable scientific input. We also thank Lincoln Stein and Linda Xiang for their assistance in the publication of the re-processed ICGC part of the PCAWG dataset. Similarly, we thank Robert Grossman, Christopher Meyer and Trevar Simmons for their assistance in the publication of the re-processed TCGA part of the PCAWG. We also like to thank Paul Wolfe and other staff of Hartwig Medical Foundation team for aligning the processing of PCAWG and Hartwig dataset.

