## Supplementary Note 1 for "Pan-cancer whole genome comparison of primary and metastatic solid tumors"

#### Cohort gathering and processing

##### Datasets

We have matched tumor-normal paired whole genome sequencing data from cancer patients from two cohorts: the Hartwig Medical Foundation (Hartwig) and the Pan-Cancer Analysis of Whole Genomes (PCAWG) cohort.

The Hartwig cohort contained patient data for which re-use for cancer research was consented by the patients and was provided under data transfer agreement (DR-247) by Hartwig Medical Foundation. The Hartwig cohort included 4,776 metastatic tumor samples from 4,460 patients. This national initiative consists of nearly 50 oncology centers inThe Netherlands and aims to improve personalized cancer treatment. To this end, Hartwig Medical Foundation sequences and characterizes the genomic landscape for a large number of patients both in clinical research projects as well as routine diagnostics (including consenting for data re-use). Furthermore, genomics data is integrated with clinical data which consists of primary tumor type, biopsy location, gender, pretreatment type before biopsy, and treatment type after biopsy. For patients with multiple biopsies taken at different timepoints, patient IDs were suffixed by a letter for the different biopsies (e.g. HMF001423A, HMF001423B). Normal samples (blood) had a mean read coverage of ~38x while tumor samples had a mean coverage of ~109x [1]. A detailed description of the consortium and the whole patient cohort has been described in detail in Priestley et al.[1]

The PCAWG cohort consisted of 2,835 patient tumors, and access for raw sequencing data for the PCAWG-US was approved by National Institutes of Health (NIH) for the dataset General Research Use in The Cancer Genome Atlas (TCGA) on 25 February 2021 under application number 100344-3 and downloaded via dbGAP download portal. Raw sequencing access to the non-US PCAWG samples was granted via the Data Access Compliance Office (DACO) Application Number DACO-1050905 on 6 October 2017 and downloaded via https://console.cancercollaboratory.org on 4 December 2017. The most recent clinical data was downloaded from the PCAWG release page (https://dcc.icgc.org/releases/PCAWG/) on Aug 2021. Normal samples (blood, adjacent tumors or distant tumors) had a mean read coverage of 39x, while tumor samples had a bimodal coverage distribution with modes at 38x and 60x. A detailed description of the consortium and the whole patient cohort has been described in detail in the PCAWG flagship paper[2].

#### Variant calling

Somatic mutation data of the Hartwig cohort were provided on 6 February 2020 and were updated on 20 October 2021. The bam files from PCAWG cohort were reformatted to FASTQ files (SamToFastq PICARD v2.1.0) which were realigned to reference genome GRCH37 lacking the GL0000XX contigs (BWA-mem v0.7.5a2[3]). These FASTQ files were used as input for the standard Hartwig analysis workflow to exclude technical noise from PCAWG and Hartwig somatic calling workflows. The Hartwig somatic calling pipeline (https://github.com/hartwigmedical/pipeline5) is hosted on a Google Cloud platform using platinum (https://github.com/hartwigmedical/platinum) that enables to run the complete Hartwig pipeline at once. A full pipeline description is explained in refs[1,4], and details and settings of all the tools can be found on their Github page (https://github.com/hartwigmedical/hmftools). Briefly, reads were mapped to GRCh37 using BWA (v0.7.17). GATK (v3.8.0) Haplotype Caller was used for calling germline variants in the reference sample. SAGE (v2.2) was used to call somatic single and multi nucleotide variants as well as indels. GRIDSS (v2.9.3)[5] was used to call simple and complex structural variants. PURPLE combines B-allele frequency (BAF) from AMBER (v3.3), read depth ratios from COBALT (v1.7), and structural variants from GRIDSS to estimate copy number profiles, variant allele frequency (VAF) and variant clonality. LINX (v1.16)[6] interprets structural variants (to identify simple and complex structural events) from PURPLE (v2.53), and also detects gene fusions, viral DNA integrations, and homozygously disrupted genes. Importantly, we ensured that mutation (simple and complex) filtering and annotation tools were run with the same versioning for PCAWG and HMF cohort.

#### Sample inclusion criteria

A selection of samples for all analyses was made based on several criteria. To exclude duplicate samples from the same patient for the Hartwig cohort, we selected the tumor sample with the most recent biopsy date, and if this information did not exist we selected the sample with the highest tumor purity. However, some patients had biopsies from different primary tumor locations (likely independent or secondary tumors). In these cases, we kept at least one sample from each primary tumor location, and when there were multiple samples from the same primary tumor location, we applied the aforementioned biopsy date and tumor purity filtering criteria. For the PCAWG cohort, we processed one tumor sample per donor and tumor sample IDs are included in Supp. Table 1 of the manuscript. As with Hartwig QC filter criteria, samples with a tumor purity lower than 20% were removed as somatic variant calling was less reliable for these samples. PCAWG samples that were gray- or blacklisted by the PCAWG consortium were also removed (see https://dcc.icgc.org/releases/PCAWG/donors_and_biospecimens). For both cohorts, we only kept samples with >=50 SNVs/indels (likely no tumor cells present in the sample), and removed an additional set of samples for several reasons including failed variant calling, insufficient informed consent for use of the WGS data, unnatural SV landscape, and one duplicate PCAWG patient (DO217844) that was also included in the Hartwig cohort. After strict QC filtering, the PCAWG whitelisted cohort includes 2,376 samples and this dataset will be made available for the cancer research community via the PCAWG resource page. The metadata for every sample including those selected for analyses is detailed in supplementary table 1. Lastly, for this study, we only selected samples from cancer types with at least 15 samples that resulted in a final dataset consisting of 3,835 Hartwig samples and 1,916 PCAWG samples.

### Impact of sequencing depth coverage and analysis pipeline on somatic variant calling sensitivity

Genomic comparisons of different WGS cancer cohorts must be conducted with caution because mutation datasets are established by distinct sequencing approaches (e.g. sequencing read depths and platforms) and somatic mutation calling workflows. The PCAWG cohort has been sequenced with an overall lower sequencing coverage (i.e. 38X and 60X) than the Hartwig cohort (109X). To assess and mitigate the impact of these confoundings, we first downsampled Hartwig samples to optimize the mapping threshold levels for the lower sequencing depths of the PCAWG cohort. Subsequently, we applied these settings to the full PCAWG cohort analyzed with the Hartwig somatic mutation pipeline. Finally, we also compared the PCAWG somatic calls obtained with Hartwig pipeline to the PCAWG consensus callset from the PCAWG resource page (https://dcc.icgc.org/releases/PCAWG/).

#### Downsampling

To validate the effect of sequencing depth on variant calling sensitivity, we selected the following 25 tumor BAM samples at random from the Hartwig cohort (109X) which we downsampled to the bimodal mean coverage depth (38X and 60X) of the PCAWG cohort using sambamba 0.6.8[7]. [HMF000054, HMF000612, HMF001536, HMF001729, HMF002822, HMF003446, HMF003796, HMF001904, HMF000318, HMF000644, HMF000920, HMF000500, HMF000774, HMF002635, HMF000164, HMF002983, HMF003913, HMF004359, HMF000458, HMF001506, HMF000097, HMF001218, HMF002379, HMF002506, HMF003755]. The matching germline samples were not downsampled because both cohorts represent similar sequencing depths for the control samples (i.e. mean depth of 39X and 38X for PCAWG and Hartwig cohort respectively). We then re-ran the identical somatic variant calling pipeline on those samples and examined variant calls of all somatic mutation types.

We first assessed the global genomic features including the ploidy level, LOH fraction and whole-genome duplication status from PURPLE. This purity ploidy estimator tool combines B-allele frequencies (BAF), read depth ratios, somatic variants and structural variants to estimate the purity and copy number profile of a tumor sample, and provides allele-specific mutation annotations (e.g. purity and ploidy corrected variant allele frequencies (VAFs)). All downsampled samples showed no difference for any of the aforementioned features which demonstrates that the calling of glocal mutation characteristics is independent of sequencing coverage with a depth of at least 38X (Fig. 1).

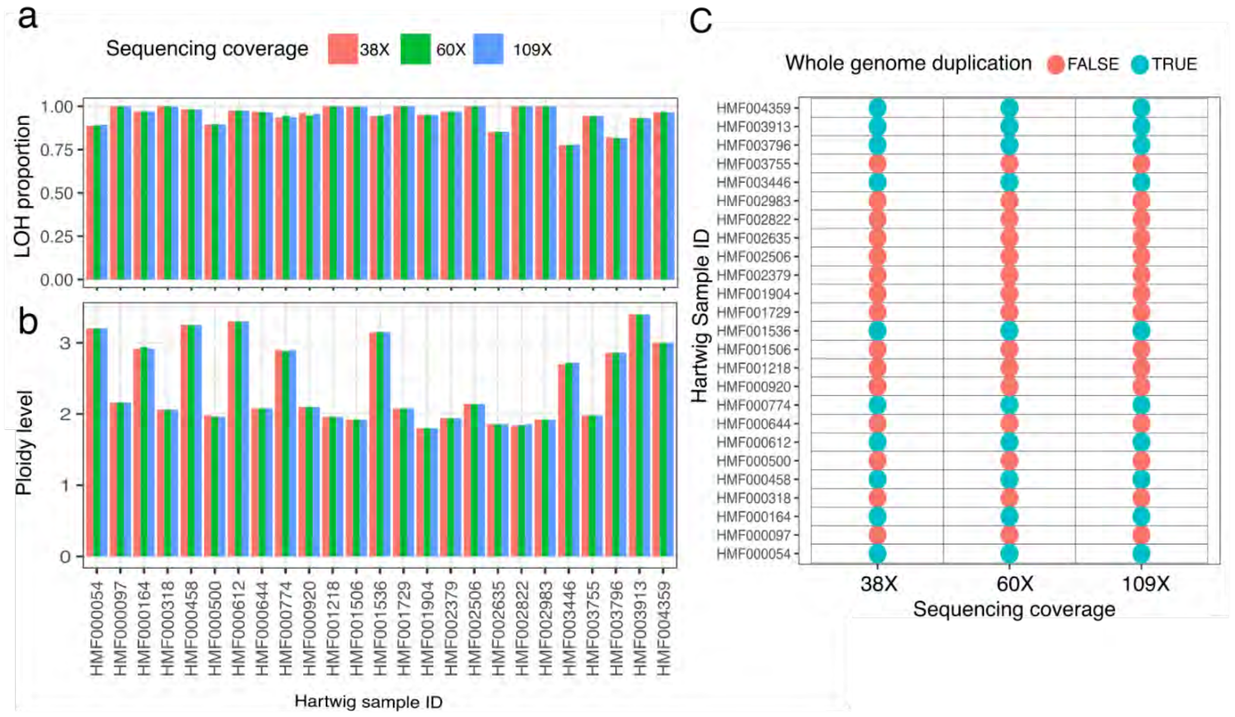


**Figure 1: Effect of sequencing depth on global genomic features.** Comparison of 25 randomly selected 109X sequenced Hartwig samples that were downsampled to 60X and 38X, the modes of the bimodal coverage distribution of the PCAWG project. The loss-of-heterozygosity (LOH) fraction **(a)**, ploidy estimates **(b)** and whole genome duplication scores **(c)** are not impacted by sequencing coverage.

We next evaluated the simple mutations of SAGE, which is the somatic mutation calling algorithm of the Hartwig pipeline. Using SAGE default settings we observed a drop in sensitivity with a fold change of 0.26x-0.04x (i.e, 26%-4%), 1.46x-0.56x and 0.35x-0.1x for SBS, DBS and Indels in 38X-60X mode, respectively. This is likely explained by the ~5 supporting read threshold within SAGE, which by default assumes a sequencing read >100x. To mitigate this effect, we increased the sensitivity for the “min_tumor_qual” flagged mutations which is the QC score for the number of high-quality reads. As such, the HOTSPOT/PANEL/HighConfidence/LowConfidence SAGE filtering settings were set to 70/70/70/90 and 70/100/100/150 for the 38X and 60X-mode respectively for “min_tumor_qual” flagged mutations. We found that these settings yielded highly similar results for the SBS, DBS and indel mutation burden for 38X and 60X (mean fold change increase of x0.00 ±0.03std.) compared to the output of the original runs with 109X depth at default settings (Fig. 2a,b).

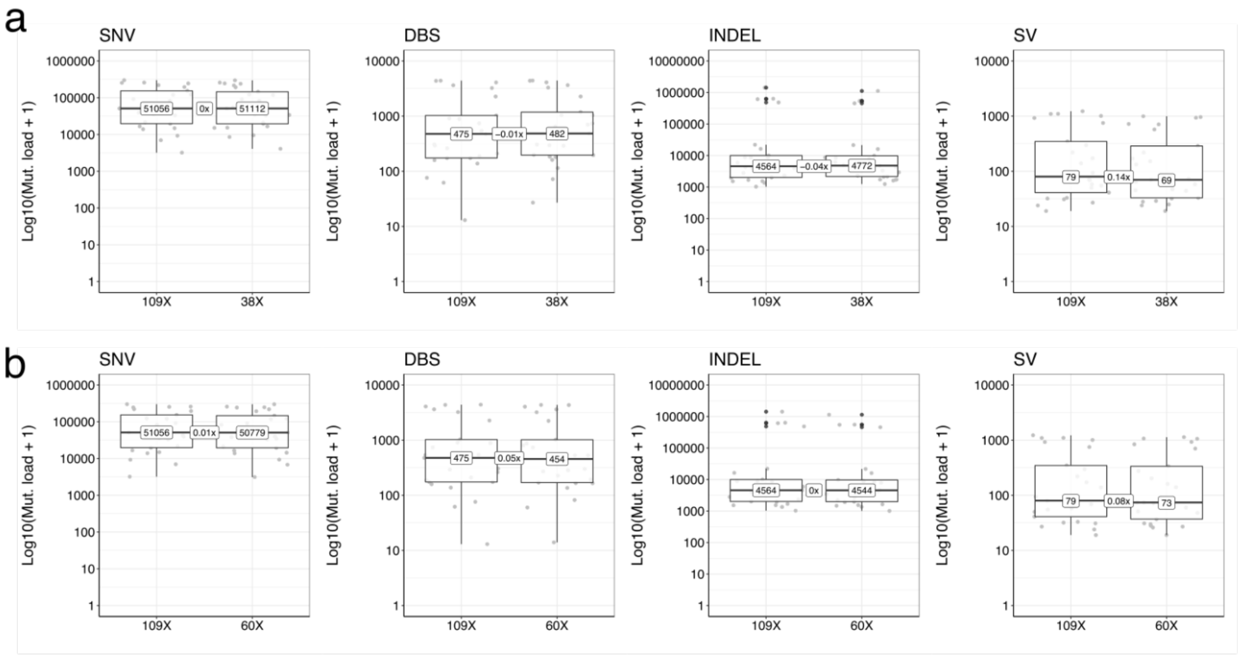


**Figure 2: Effect of sequencing depth on somatic mutations.** Tumor mutation burden comparison of 25 randomly selected 109X sequenced Hartwig samples that were downsampled to 38X **(a)** and 60X **(b)** for SNV, DBS, INDEL and SV mutation types. The boxplots show the median mutation burden for each sequencing depth mode as well as the median fold change sensitivity loss for every mutation type.

We also observed a minimal effect on the driver landscape. For the 172 reported drivers in the 25 Hartwig samples sequenced at 109X, 95% and 94% of the drivers were recalled in the 38X and 60X-mode respectively. Given the consistent results in passenger and driver mutation detection over the tested sequencing depth modes, we have applied these two SAGE filter modes to all PCAWG samples depending on their respective sequencing depth bucket to normalize the sensitivity impact on mutation calling between the PCAWG and Hartwig cohort.

Next to simple somatic mutations, we also validated structural variants from LINX[6]. This tool integrates copy number profiles and the SV calls from PURPLE[1] and GRIDSS[5] that enables the clustering and chaining of genomic rearrangements. To evaluate the impact of sequencing depth on SV calling, we analyzed the 60X and 38X downsampled samples and observed a decrease in sensitivity with a fold change of 0.08X and 0.14X (i.e., 8%-14%), respectively (Fig. 2a,b). Given the complexity of the integrated SV calling workflow, we have not adjusted threshold settings for the SVs. As a result, particularly samples sequenced with ~38X have a decreased sensitivity to call SVs. However, only a few PCAWG cancer type cohorts consist of a high proportion of 38X samples (Fig. 3a). To estimate the effect of sequencing depth on cancer type level for the PCAWG cohort, we have assessed the sensitivity loss at varying proportions of 38X and 60X sequenced samples per cancer type for all mutation types.

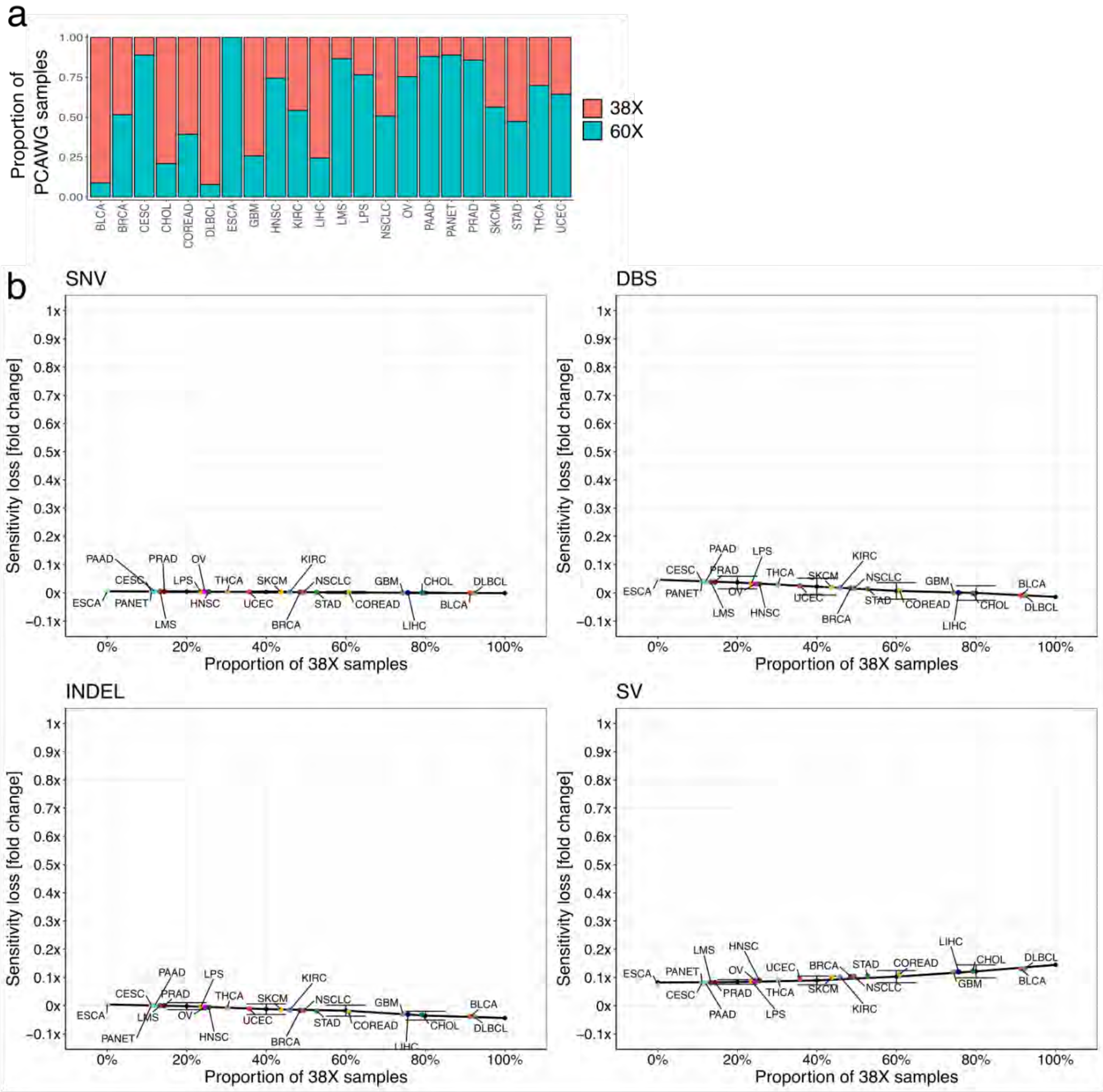


**Figure 3: Effect of sequencing depth on PCAWG cancer type. (a)** Every PCAWG cancer type cohort consists of a different proportion of 38X and 60X sequenced samples. **(b)** To reveal the sensitivity loss across all cancer types for the SNV, DBS, INDEL and SV counts, we performed a bootstrap approach. Here, we randomly assigned 38X and 60X samples in line with each proportion bin and computed the median fold change difference. We performed this procedure 100 times and calculated the average (±StDev) of the fold changes across all bootstraps. Subsequently, the obtained fold change scores were fitted using a linear regression model that allowed us to predict the sensitivity loss for every 38X/60X proportion. The sensitivity loss for every cancer type of every mutation type is depicted by coloured dots.

As expected, the fold change is negligible for SBS (ranging from -0.007X to 0.005X), DBS (ranging from -0.01X to 0.04X) and indels (ranging from -0.04X to 0.007X) for diffuse b-cell lymphoma (DLBCL, 92% of samples sequenced at 38X) and esophagus cancer (ESCA, fully consisting of 60X sequenced samples), respectively (Fig. 3b). The SVs comprise the largest impact on sequencing depth with a fold change ranging between 0.13X for DLBCL and 0.07X for ESCA.

However, these lower SV detection sensitivities only marginally explain the observed increases in SV burden in the metastatic Hartwig tumors as compared to the primary PCAWG tumors. In fact, the SV burden statistics remained unchanged after applying the tumor type-specific correction factors (see Fig. 4b). Among the 14 significant cancer types with increased SV burden, only breast (BRCA), NSCLC and cervix cancer (CESC) lost significance after applying this correction factor. In fact, in CESC the fold change almost did not change after correction (fold change of x2.6 raw versus x2.4 adjusted) and the loss of significance is more likely related to the high variation in SV burden between samples in the primary and metastatic cancer cohort as well as by limited sample size.
Finally, the mutation burden for SNV, DBS and INDEL are almost identical between the raw and the adjusted burden (Fig. 4a). Consequently, the observed fold changes between primary and metastatic are highly consistent between raw and adjusted mutation counts. Overall, this downsampling analysis demonstrates that, after minimal adjustments, the lower sequencing coverage of PCAWG samples marginally impacts the mutation calling sensitivity for all mutation types.

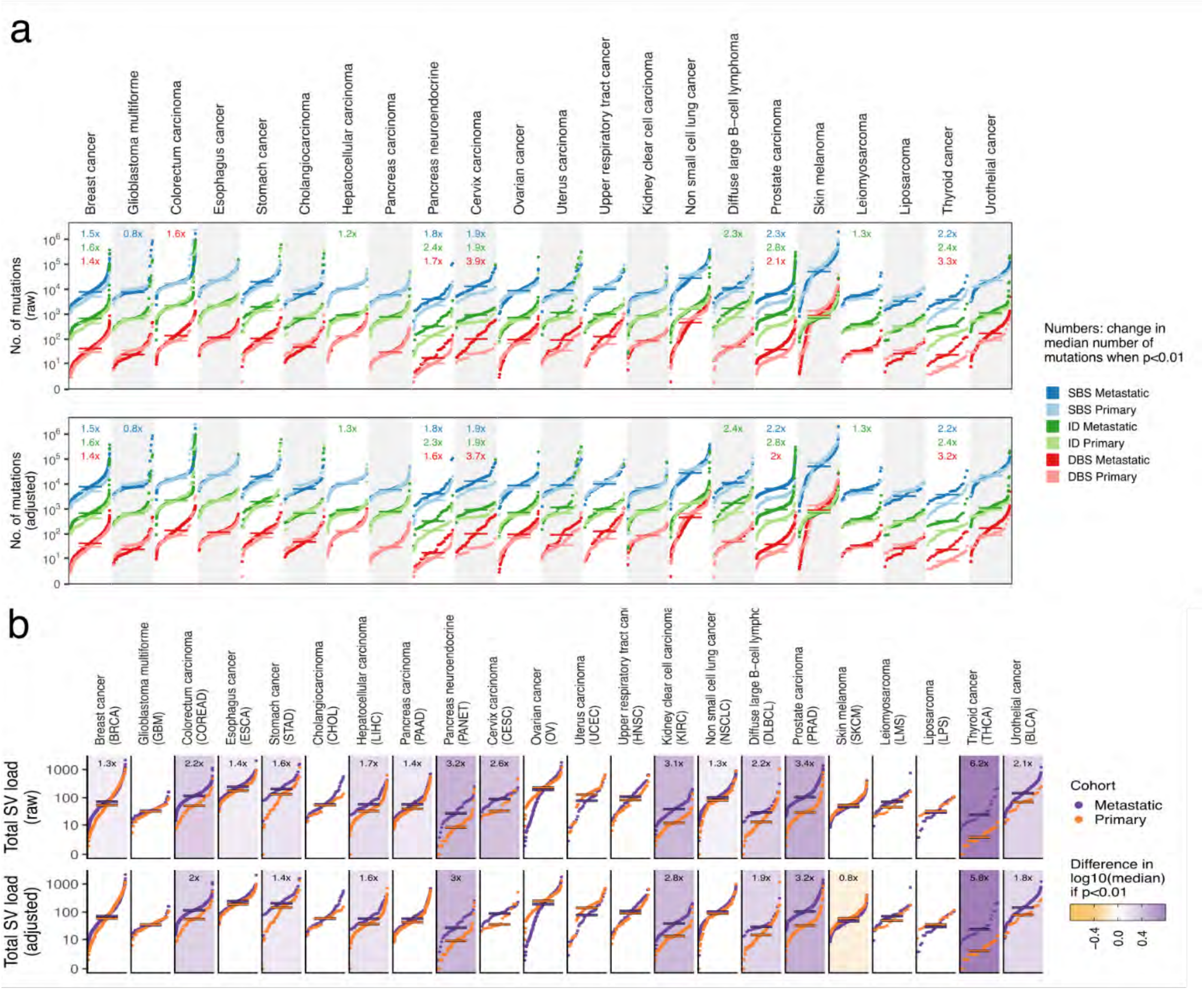


**Figure 4: Effect of sequencing depth on mutation burden.** Cumulative distribution function plot (samples were ranked independently for each variant type) of tumor mutation burden for each cancer type for SMNVs **(a)** and SVs **(b)**. The SMNV plot **(a)** shows on top the cumulative distribution function plot for each tumor type for SBS (blue), INDELs (green) and DBS (red). Below, the cumulative mutation distributions adjusted for the estimated sensitivity loss per cancer type caused by differences in 38X/60X sequencing depth proportion. Similarly, on top of the SV plot **(b)** the cumulative distribution function plot for each tumor type for the SV mutation burden and below the cumulative mutation distributions adjusted for the estimated sensitivity loss per cancer type caused by differences in 38X/60X sequencing depth proportion. Horizontal lines represent median values. Fold change labels are included only when Mann-whittney p-value < 0.01.

#### The impact of clonality on variant calling

We found that metastatic lesions generally present lower intra-tumor heterogeneity compared to primary tumors as a result of the severe evolutionary constraints imposed by anti-cancer therapies and/or the metastatic seeding event. The clonal outgrowth of subclonal mutations prior to the evolutionary process will become clonal (i.e., “clonality ilusion”) and more easily detectable by bulk WGS, which may in part explain the increased TMB and SV burden in metastatic samples of some cancer types. To assess the impact of clonality on SNMV and SV mutation burden, we compared primary and metastatic samples independent of clonality. For this, we computed the average proportion of clonal mutations by dividing the number of clonal mutations by the total mutation burden (see methods). We selected samples with high global clonality (defined as those with a fraction of >85% clonal mutations) across primary and metastatic tumors, and created sub-cohorts populated by these samples. This approach resulted in a sufficient number of representatives to compare both stages of tumor development independent of clonality.

As depicted in Figure 5 (see below), the fold change differences for SBS, DBS and indel mutation burdens of clonal primary and metastatic samples (Figure 5a) were very much in line with the mutation burdens not filtered by clonality (main Figure 2). This was most obvious for breast, cervix and prostate carcinoma where we noticed highly similar fold change rates for all mutation types. Thyroid and pancreas neuroendocrine cancers lost their significant increase, but this effect can be attributed to the reduction of the sample size. Indeed, the SBS, DBS and indel fold change metastatic increase was 1.6x, 1.9x and 1.7x, respectively, for clonal pancreas carcinoma which is in good agreement with the cohort not filtered on clonality (i.e., 1.8x, 2.3x and 1.7x respectively - main Figure 2). Similarly, clonal cervix cancers showed a fold change increase of 2.1x, 1.9x and 4.1x, respectively, whereas we found a very similar increase of 1.9x, 1.9x and 3.9x for the unfiltered cervix cohort (main Figure 2). Performing mutational signature analysis on the clonal mutations revealed a contribution landscape that showed a strong resemblance to the mutations contributions of the unfiltered samples (Figure 5a, bottom panels).

Similar conclusions can be drawn from the SV comparison. After subsetting for clonal samples, we found that 9 metastatic cancer types remained their significant impact on SV burden increase (Figure 5b). As with SNMVs, the reduced sample size explained the loss of significance in the 5 cancer types which encompassed a significant metastatic SV burden increase in the unfiltered cohort. This is shown by the highly similar fold change differences between clonal and unfiltered cancer cohorts: breast (1.2x- clonal versus 1.3x - unfiltered), esophagus (1.3x vs 1.4x), cervix (2.4x vs 2.6x), non small cell lung (1.2x vs 1.3) carcinoma. Only clonal metastatic pancreas neuroendocrine cancers showed a slightly lower SV burden (2.2x) compared to metastatic pancreas neuroendocrine cancer samples not filtered by clonality (3.2x).

In conclusion, this analysis demonstrates that cancer samples with comparable fraction of clonal mutations show highly similar SNMV and SV mutation profiles to those cancer samples not filtered on clonality. Therefore, the clonal selection/expansion imposed by the evolutionary bottlenecks seems to only marginally contribute to the increased SNMV and SV mutation burdens in metastatic cancers as described in main Figure 2 and main Figure 4.

Supp. Note 1 of *“Pan-cancer whole genome comparison of primary and metastatic solid tumors"*

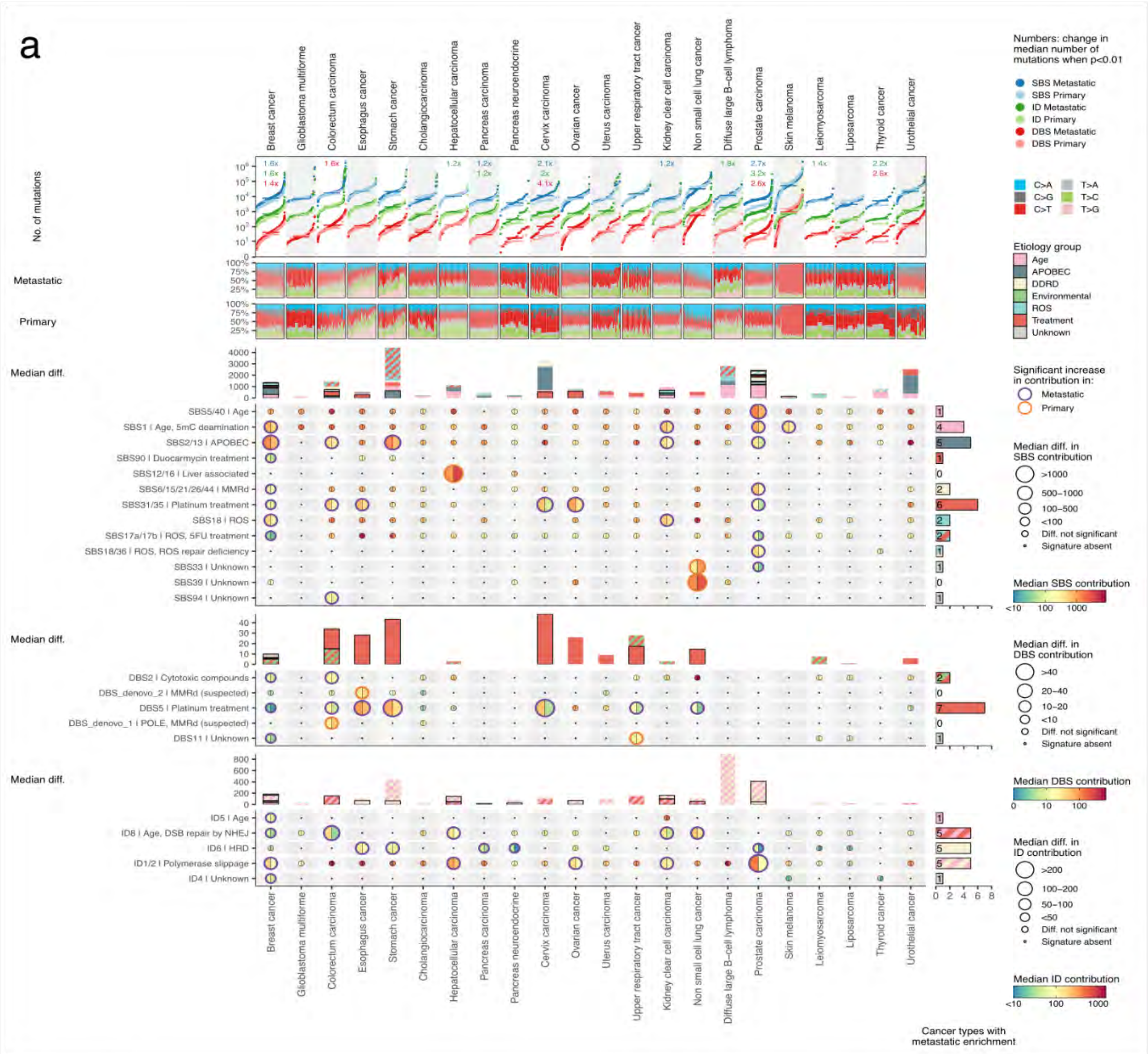


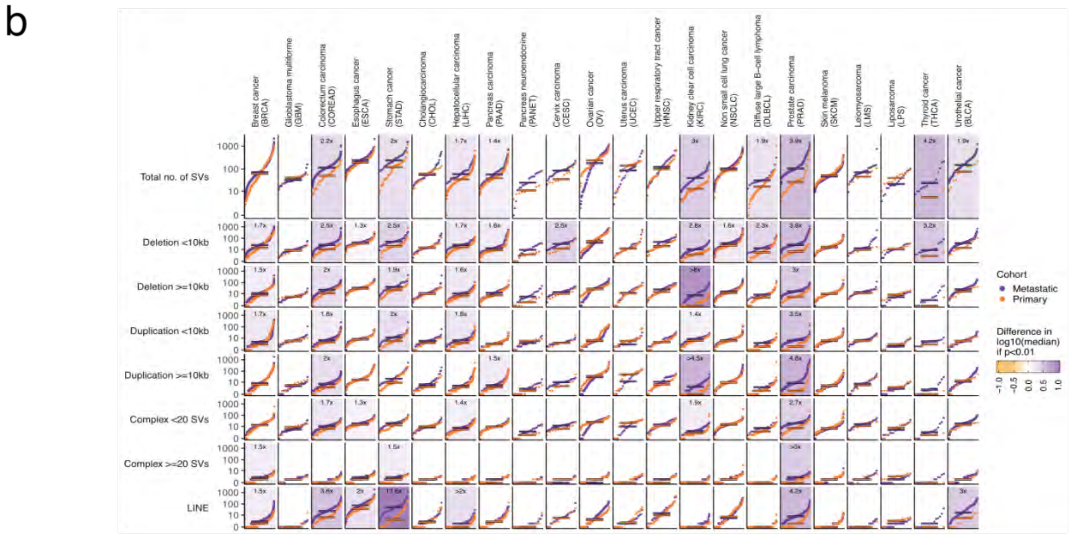

**Figure 5: Effect of clonality on passenger mutation landscape.** Each cancer type cohort was subsetted with samples having a clonality of at least 0.85. The SMNV plot **(a)** shows on top the cumulative distribution function plot for each tumor type for SBS (blue), INDELs (green) and DBS (red). Below, the mutational signature contribution landscape for samples with a clonality estimate of at least 0.85. **(b)** The cumulative distribution function plot for each tumor type for the SV mutation burden for samples with a clonality of at least 0.85. Below the distribution plots stratified by SV type. Horizontal lines represent median values. Fold change labels are included only when Mann-whittney p-value < 0.01.

#### The impact of pipeline differences on variant calling

Next, we compared the genomic features obtained with the Hartwig pipeline to the consensus somatic calls that were generated by the PCAWG Working Groups available on the PCAWG resource page. Many mutational features are closely connected with correct sample purity determination and ploidy level of the sample. We therefore assessed the WGS-based purity estimates between PCAWG consensus calls and the ones obtained from the Hartwig pipeline. We found that for 87% of the whitelisted samples of the ICGC part (n=1,916) the purity estimates were within the 10% range difference between the two pipelines (Fig. 6). Only 12 samples (0.6% of the samples) showed a remarkable difference where Hartwig estimated the purity close to 100% whereas PCAWG reports a purity level lower than 50%. Although improvements on purity estimations have been made with the latest version of purple, the low number of discrepant samples can be explained by the low levels of aneuploidy for these samples. For the ploidy levels, we observed that 90.0% of the samples showed a similar ploidy estimate (i.e. within a 10% range difference of ploidy level) between the two pipelines (Fig. 6). The remaining samples show either a doubling (4.2%) or a halving (3.4%) of the ploidy scores. Finally, 93.4% of the samples shared the same whole-genome duplication status.

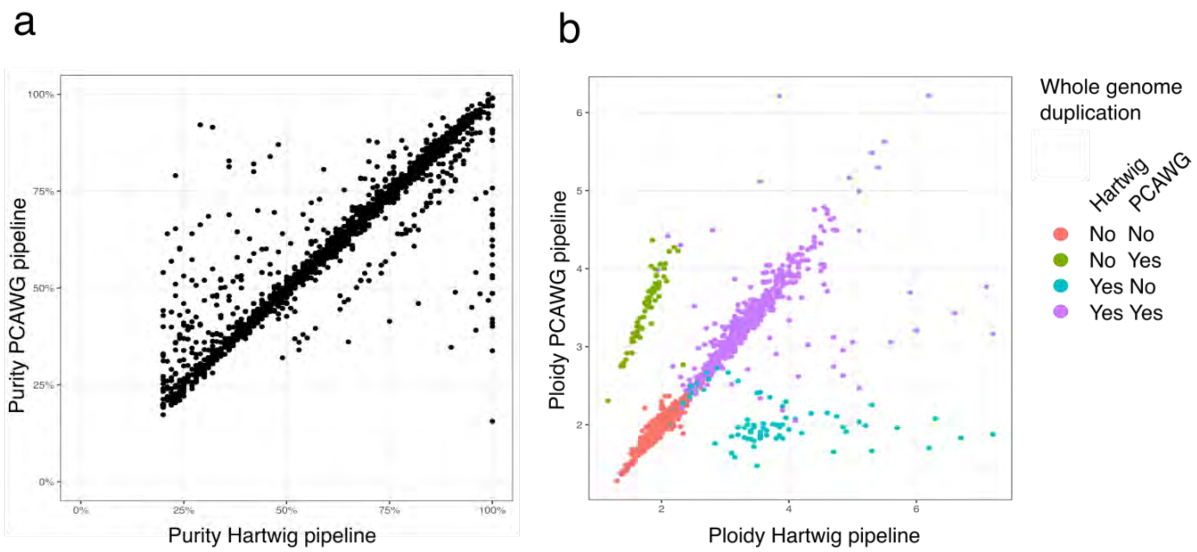


**Figure 6: Effect of mutation calling pipeline differences for purity and ploidy scores.** Comparison of all whitelisted ICGC-PCAWG samples for purity **(a)** and ploidy **(b)** obtained by the Hartwig pipeline to the scores provided by PCAWG resource page. The purity estimates were within the 10% range difference between the two pipelines for 87% of the samples. 90.0% of the samples showed a ploidy estimate within the 10% range difference and most of the remaining samples show either a doubling (4.2%) or a halving (3.4%) of the ploidy scores and explain most of the whole genome duplication estimates.

To compare the mutational burden for each mutation type between PCAWG consensus callset and mutations obtained with Hartwig pipeline, we excluded the trinucleotide MNVs as well as SV calls with less than 500 base pairs in length because these are not called by the PCAWG pipeline. The figure below shows that both pipelines share similar calling sensitivity for SBS as shown by the median fold change of 0.08x (Fig. 7). The DBS (fold change of 0.36x) and indels (fold change of 0.34x) are more sensitively called by the Hartwig pipeline. SAGE includes the local phasing information for DBS and indel mutations (co-existence on the same read) and integrates a context-specific scoring system for indels that is particularly applied to indels in repeat contact. These SAGE features may explain the higher calling sensitivity for DBS and indel mutation types. We also observed that the mutation spectra were overall in good agreement (i.e. cosine similarity > 0.90 for 96%, 80% and 94% for SNV, DBS and indel respectively) between both pipelines (Fig. 7). As expected, the largest discrepancies were observed for samples with low mutation burden.

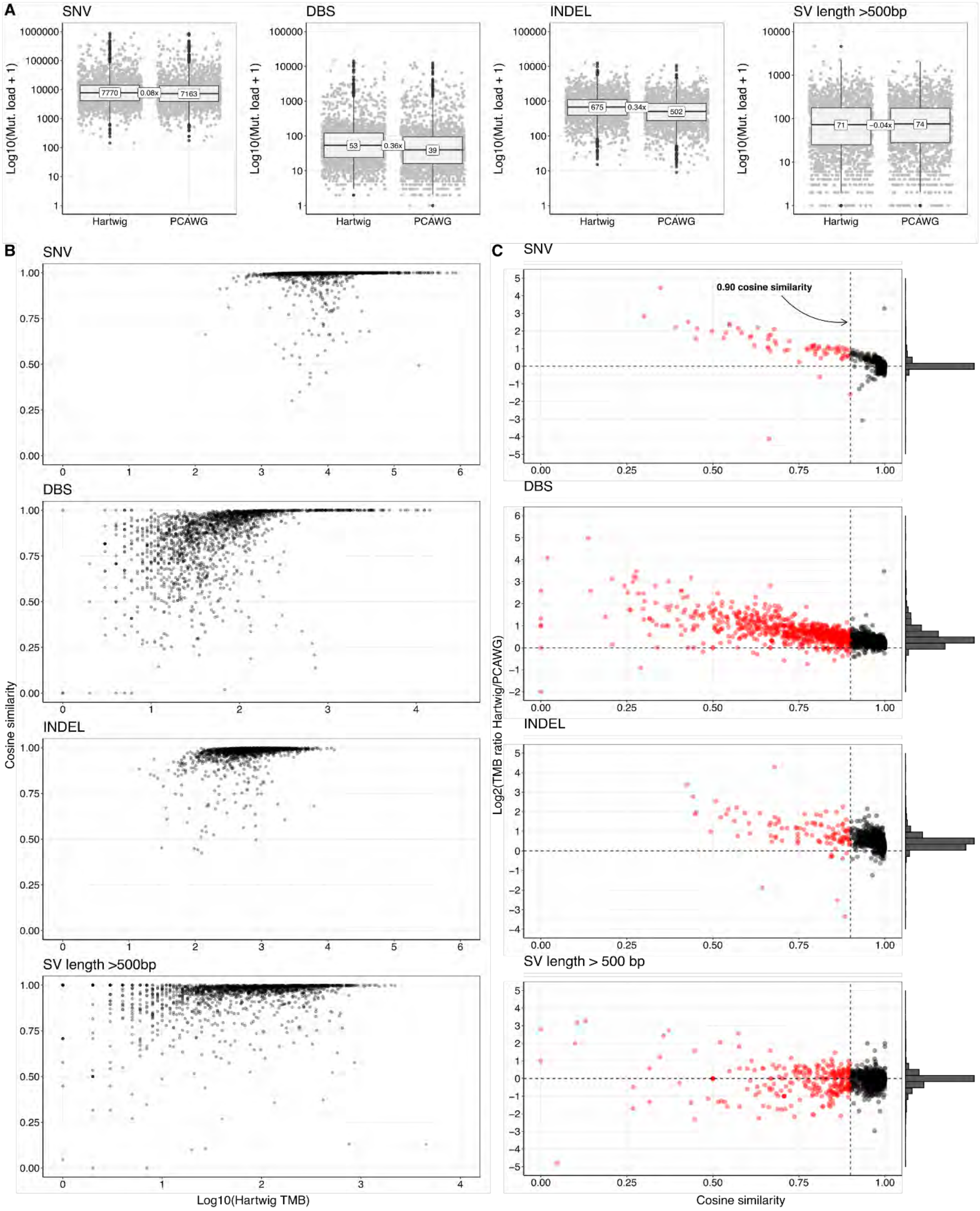


**Figure 7: Effect of mutation calling pipeline differences for somatic mutations. (a)** Comparison of all whitelisted ICGC-PCAWG samples for SNV, DBS, INDEL and SV counts obtained by the Hartwig pipeline to the mutation counts provided by PCAWG resource page. The PCAWG consensus calls show a small difference in sensitivity for DBSs,

indels, and nearly the same sensitivity for SNVs and SVs. The fold changes of all mutation types for the two sequencing depth modes are depicted in table 1. **(b)** Most of the samples harbor a highly similar mutation context (cos sim > 0.90). **(c)** Only samples with a very low mutation burden show a different mutation context.

A comparison in driver landscape between Hartwig and PCAWG pipeline is more challenging because the Hartwig and PCAWG driver analysis rely on different driver detection tools and driver reference databases. Nevertheless, we observed that 90% of the samples showed a similar number of driver hits per sample between PCAWG driver consensus callset (only ICGC part; n=1613) and drivers obtained with the Hartwig pipeline. 32% of the samples showed the same number of driver genes per sample, while 41% and 17% of the samples showed a difference of respectively 1 and 2 in driver hits per sample. On gene basis per sample, we found that ~50% of the 4738 driver genes reported by either the Hartwig or PCAWG pipeline were found by both pipelines, while ~34% were Hartwig-unique and ~16% PCAWG unique. This may be explained by the inclusion of cancer driver databases in the Hartwig driver analysis, while the PCAWG driver catalog was established with a compendium of *de novo* driver discovery tools. Indeed, a higher similarity in gene drivers was found for the top-20 mutated genes with 68% overlap in gene driver landscape between PCAWG and Hartwig pipeline and ~25% Hartwig-unique and 7% PCAWG-unique. Taken together, the PCAWG mutation callset generated by Hartwig analytical pipeline is in line with the PCAWG consensus mutation callset. The ICGC part of the Hartwig processed PCAWG dataset is made available as a WGS cancer genomic resource at the PCAWG release page (https://dcc.icgc.org/releases/PCAWG/Hartwig).
